# Laboratory evolution enhances resilience of a symbiont yeast and its honeybee host against agrochemical exposure

**DOI:** 10.64898/2026.07.10.737698

**Authors:** Simone Mozzachiodi, Flavia Di Cesare, Stephan Kamrad, Xiaoteng Jiang, David Scheidweiler, Jack Rogan, Sandip Kumar Patel, Anna Lindell, Luisa Faria, David Baracchi, Kiran R. Patil

## Abstract

Yeasts are key microbial members of several ecosystems. Yet, the impact of widespread chemical pollutants on yeasts is only sparsely studied. Here we report the effect of >1000 chemical pollutants on fourteen diverse yeast species spanning the Saccharomycotina subphylum. *Starmerella bombicola*, a symbiont of various bee species, was the most sensitive and inhibited by several fungicides as well as by non-fungicides. To identify the molecular basis of this ultra-sensitivity, we selected resistant lineages against nine chemicals using adaptive laboratory evolution. Whole-genome-sequencing uncovered convergent evolution on *YBP1*, a key regulator of oxidative stress. Proteomic analysis confirmed the protective role of oxidative stress response pathways, including proteins encoded by horizontally transferred bacterial genes. We find that the evolved *S. bombicola* stably colonized the bee gut and ameliorated the negative effect of paclobutrazol, a plant hormone regulator, on gut microbes, sucrose responsiveness, and learning. Our findings demonstrate how laboratory evolution can be used to mitigate the negative impact of chemical pollutants on pollinators.

---

Environmental pollution by human-made chemicals has now exceeded the safe planetary boundary with far-reaching implications across ecosystems^1,2^. Eco-toxicological studies on multicellular organisms such as humans^3^, fishes, and insects^4^ are mapping the impact of pollutants on multi-cellular biological systems. Recent studies have also indicated negative impact of chemical pollution on microbial ecosystems^5,6^. However, the impact of chemical pollutants on environmental and host-associated microbes is only sparsely studied. This knowledge-gap poses a major hurdle for mechanistic understanding of eco-toxicological effects and thereby hinders development of strategies to enhance host resilience through their microbiomes.

Various studies have shown that certain species of yeasts and fungi are vulnerable to specific agrochemicals (e.g.^7,8^). While systematic studies investigating chemical toxicity in host-associated bacteria are emerging^9,10^, there is a lack of studies spanning both phylogenetic and chemical diversity of yeasts and fungi despite their key role in host-microbiome interactions^11,12^ and ecosystems health in general (Bahram & Netherway, 2022). In this study, we assessed toxicity of over 1000 common chemical pollutants on 14 fungal species that are found across diverse ecological niches, ranging from wine fermentation to human and honeybee microbiomes ^14^. Mechanistically, we focus on the yeast *Starmerella bombicola*, which exhibited sensitivity to many chemicals in our study. The genus *Starmerella* is associated with several wild as well as managed bee species worldwide^15,16,17,18^. With circa 30% of the total yeast counts in healthy honey-bee guts, *Starmerella* is a common bee gut symbiont. It is amongst the most abundant yeast genus in the honeybee microbiomes^19^ and positive physiological effects have been described in bumble bees such increasing offspring survival^17^. We investigate the molecular basis of *S. bombicola*’s chemical sensitivity by using adaptive laboratory evolution coupled with genomic and proteomic analyses and demonstrate how laboratory-evolved symbionts can improve the resilience of honeybees.

## Results

### Agrochemicals including non-fungicides inhibit diverse yeast species

To systematically assess the susceptibility of yeasts to commonly used chemical contaminants, we used a comprehensive library of 1025 chemical pollutants^9^ (**Supplementary Figure 1a**). Most of these (829 of 1025, 80.9%) are active ingredients in the current or previously marketed pesticide formulations and agricultural chemicals. The library also contains 119 known pesticide metabolites, biotransformation or degradation products with potential toxic effects^20^.To span yeasts of ecological importance, we selected 14 species with diverse ecological niches (**Fig 1a-b, Supplementary Table 1**). These species cover the entire breadth of the *Saccharomycotina* subphylum and includes human pathogens such as *Candida albicans*, widely used biotechnological yeasts, such as *Saccharomyces cerevisiae* and *Komagataella pastoris,* and species associated with pollinators, such as *Starmerella bombicola*.

**Figure 1.**
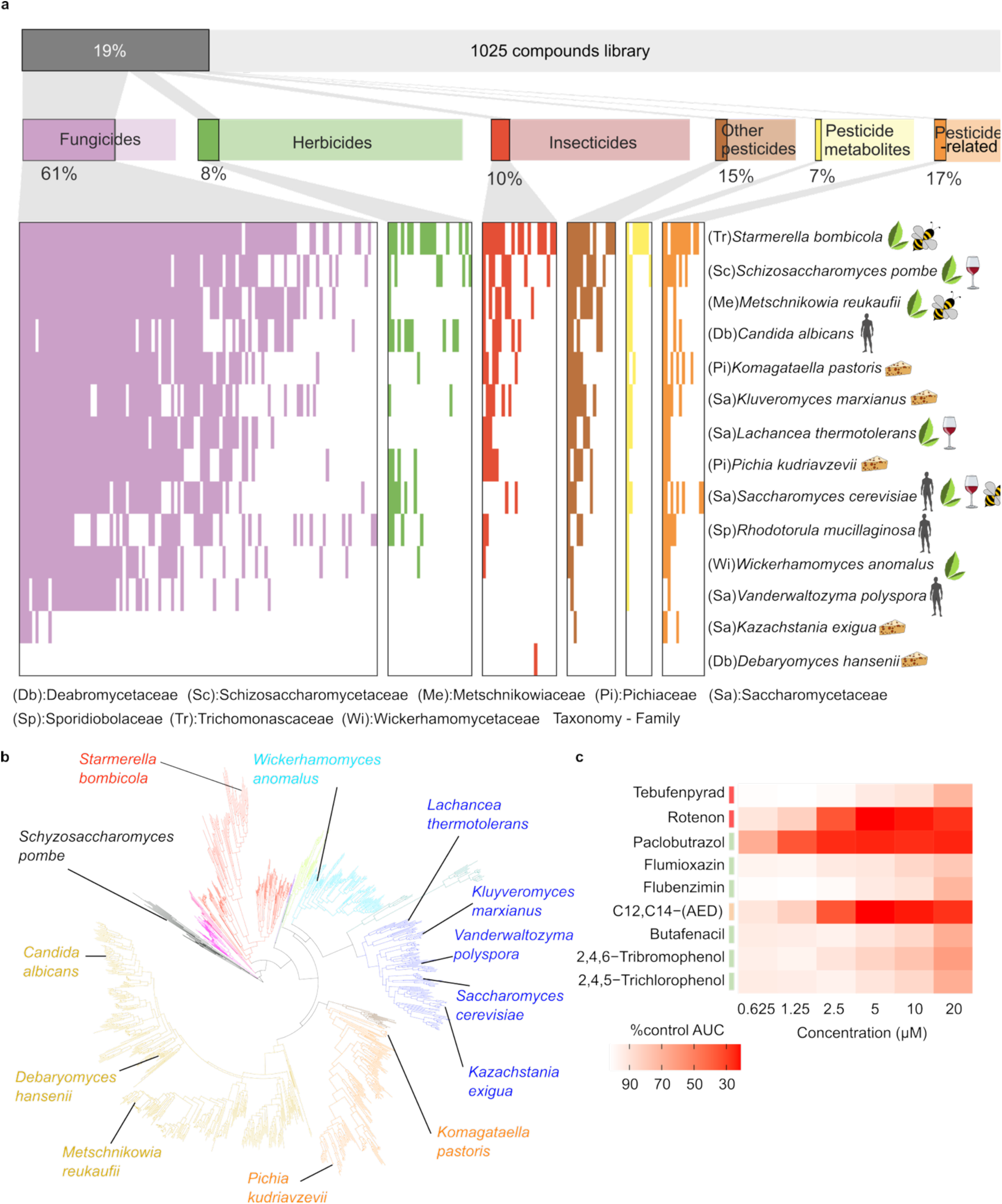
Widespread off-target toxicity of pesticides across different yeast species. **a.** Overview of the agrochemicals impacting the growth of 15 diverse yeast species. Filled bars in the heatmap show significant interactions between chemicals (columns) and yeast species (rows). Yeast growth is measured as area under the curve (AUC, Methods). Interactions are deemed significant if p<0.05 (two-sided Z-test, Bonferroni correction, n=3 biological replicates) and at least 15% reduction in growth. Two letter code in parentheses denote the taxonomic family to which the species belongs. Ecological origins/habitats are depicted as pictograms. **b.** Phylogenetic tree of Saccharomycotina subphylum marking the position of the tested yeast species. The tree is based on genomic data from^14^. The branches are coloured according to the taxonomical order. **c.** Heatmap representing the concentration dependent decrease of *S. bombicola* growth across a series of selected chemicals. The chemicals are coloured according to their class as in **a**.

The 14 yeast species were then tested against 1025 chemicals for their impact on growth using liquid culture growth in 96 well plates (14350 chemical-yeast interactions in total, **Supplementary Table 15**). All chemicals were used at a fixed concentration of 20 µM, which is at the lower range of typical field applications (Gandara et al., 2024). Cells were grown in a rich medium, YPD, which is a NCYC (National Collection of Yeast Cultures, UK) recommended media and supported the growth of all tested species (**methods**). This enabled systematic comparison across the yeast phylogenetic breadth. Three biological replicates executed on separate days were performed for all chemical-yeast pairs. Low median (across species) coefficient of variation, 5.4±3%, low fraction of unexplained variance (FUV) of 14.7±14% (**Supplementary Figure 1b**), and good agreement between replicates (**Supplementary Figure 1c**) confirmed the robustness of our screen. We uncovered 1027 growth inhibitory interactions between pollutants and yeasts. Over 19% of the tested chemicals (194/1025) inhibited at least one yeast species (>15% reduction in growth in at least 2 out of 3 biological replicates, p_adj_ <0.05, Bonferroni-corrected, two-sided Z-test). Notably, 84 of the yeast inhibiting compounds are not classified as fungicides suggesting extensive off-target toxicity of pesticides (**Fig 1a**).

Amongst the 14 tested yeasts, the bee gut symbiont *Starmerella bombicola*, a species branching close to the root of the Saccharomycotina subphylum (**Fig 1b**), exhibited the highest sensitivity. It was inhibited by 150 compounds, 57 of which are not classified as fungicides. Phylogenetically close to *S. bombicola*, *Schyzzosaccharomyces pombe* was the second species per number of hits, inhibited by 110 compounds while the more distant *Candida albicans* was the second species per hits on herbicides confirming that phylogenetic distance and species sensitivity does not clearly correlate. *Saccharomyces cerevisiae* was not particularly affected from insecticides and herbicides while other food associated yeasts such as *Kazachstania exigua* (n=9 hits) and *Debaryomyces hansenii* with 1 hit showed the lowest sensitivity across our panel. To further characterize the *S. bombicola* sensitivity, we tested 9 compounds at 6 concentrations (**Fig 1c**). The deleterious effects on growth were found at concentrations as low as 0.625 µM, well below that of concentrations found in agricultural settings.

Overall, our comprehensive screen shows that off-target toxicity to chemical pollutants is common in yeasts across different ecological and phylogenetic ranges and that *S. bombicola* is particularly affected by fungicides and non-fungicides alike.

### *Starmerella bombicola* adapts to pesticides with emergent pleiotropy

*Starmerella bombicola* exhibited the broadest sensitivity to agrochemicals. Given its ecological relevance as a bee gut symbiont, we set out to uncover the molecular mechanisms underlying its broad sensitivity using adaptive laboratory evolution (ALE), which can provide unbiased mechanistic insights into molecular pathways involved in the sensitivity^21,22^.

For our experimental evolution approach, we selected 9 compounds including 3 insecticides (Flubenzimin, Rotenone, Tebufenpyrad), 3 herbicides (Butafenacil, Flumioxazin, Paclobutrazol) and 3 other chemical pollutants (Tribromophenol, 2,4,5-Trichlorophenol, Alkyl(ethylbenzyl)-dimethylammoni) (**Supplementary Table 2**). Concentration for each compound was set corresponding to pre-evolution growth reduction between 20 and 50%, which spanned the range between 1.25 µM (Paclobutrazol) to 80 µM (Rotenone). These concentrations are well in the range of ecological observations^23^.

For each chemical, we evolved 4 independent populations over 20 growth and dilute cycles (**Fig 2a**). In 6 cases, we observed rapid adaptation dynamics with increasing optical density (OD), a proxy of culture biomass. In the remaining 3 cases (Flubenzimin, 2,4,5-trichlorophenol and Alkyl(ethylbenzyl)-dimethyl ammonium) all replicate populations failed to grow after the first five passages (**Figure 2b, c**). Thus, while the *S. bombicola* has potential to adapt to chemical stressors, it is not universal and may be contingent on the underlying toxicity mechanism.

**Figure 2.**
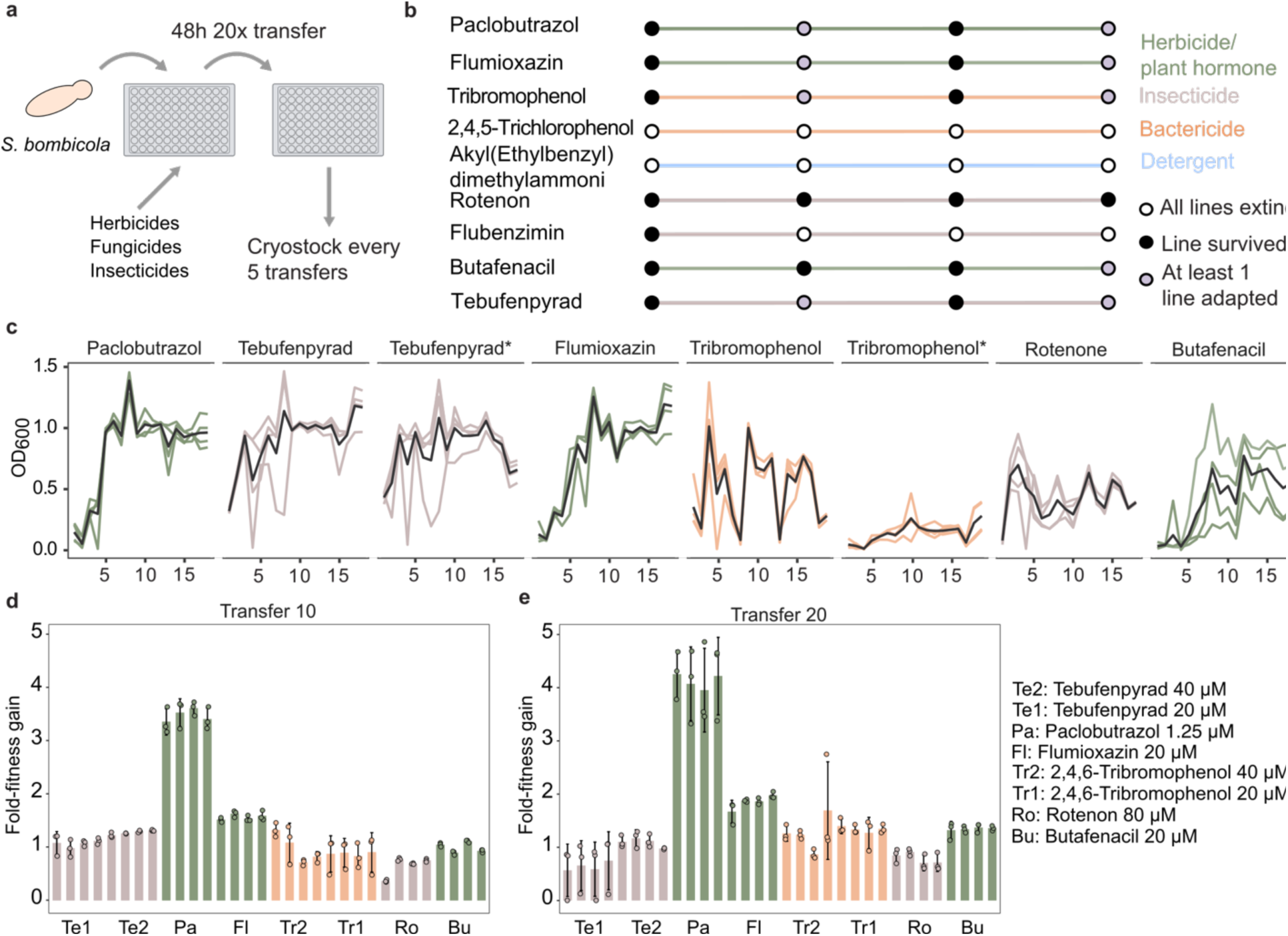
Adaptative laboratory evolution of *Starmerella bombicola.* **a.** Experimental design of the adaptive laboratory evolution experiment. **b.** Survival outcomes of the evolution experiment for each chemical pollutant. Each point represents 5 transfers and the status of the populations at each of the 5 transfers is colour coded. Lineages at the 10th and 20th transfers were phenotyped. Adapted status call required at least one of the 4 lineages showing improved growth compared to the control. **c.** Lines depict the OD variation relative to the YPD controls during 18 transfers for all the populations that reached the end of the experiment. The y-axis reports the optical density after blank subtraction and controls normalisation. Data in **Supplementary Table 14**. Stars highlight compounds used at two different concentrations. The black line shows the average OD across the 4 lineages (evolving in parallel). **d.** Bar-plot reporting the increase in fitness after 10 transfers measured as ratio of growth (AUC, Area Under Curve) between evolved and the ancestral strain. n=3 biological replicates and 4 independent populations. Fitness-gain of 1 indicates no appreciable increase in the fitness. The error bars represent the standard deviation while the bar height is the mean. **e**. As in **d** but after 20 transfers of the evolving populations.

We next compared the growth of the adapted populations after 10 and 20 transfers to the parental strain for 5 chemicals (Butafenacil, Rotenone, Flumioxazin, Paclobutrazol and Tebufenpyrad). In 2 cases, adaptation occurred after 10 transfers while by the end of the experiment, adapted populations were evident in all 5 conditions (**Figure 2d, e**). The populations evolved in Paclobutrazol, a chemical used in treatment of flowering plants, to the highest fitness gain with an average four-fold improvement (**Figure 2e**). To test the stability of the observed phenotype, we isolated single clones (n=24) from one of the populations evolved in Paclobutrazol. These single clones replicated the improved fitness observed for the evolved populations suggesting strong selection during the laboratory evolution (**Supplementary Figure 2**). Interestingly, the isolates adapted to Paclobutrazol showed cross-protection against Flumioxazin, an insecticide, and Butafenacil, a common herbicide used together with glyphosate hinting at pleiotropic resistance against chemicals with similar toxicity mechanisms (**Supplementary Figure 2**).

### Genomic mutations identify *YBP1* as a molecular target of convergent evolution

To understand the genetic basis underlying the adaptation of *S. bombicola*, we performed whole-genome-sequencing (WGS) of the populations evolved under 5 chemicals for which we confirmed adaptation (Paclobutrazol, Tebufenpyrad, Butafenacil, Flumioxazin and Tribromophenol). As a control, we sequenced 4 populations subjected to the same evolution conditions except for the chemical stressors. Genomics analysis showed no chromosomal aneuploidy but identified 45 unique mutations in protein-coding regions of the adapted populations, of which 42 are single nucleotide mutations and 3 indels spanning 33 genes (**Fig 3a left**). 14 genes with single point mutations shared an ortholog with other *Saccharomycetaceae* and in *Saccharomyces cerevisiae* (**Supplementary table 3**). For the other 19 mutations, no orthologs were found suggesting selection on non-syntenic genes private to *S. bombicola* or the *Starmerella* genus.

**Figure 3.**
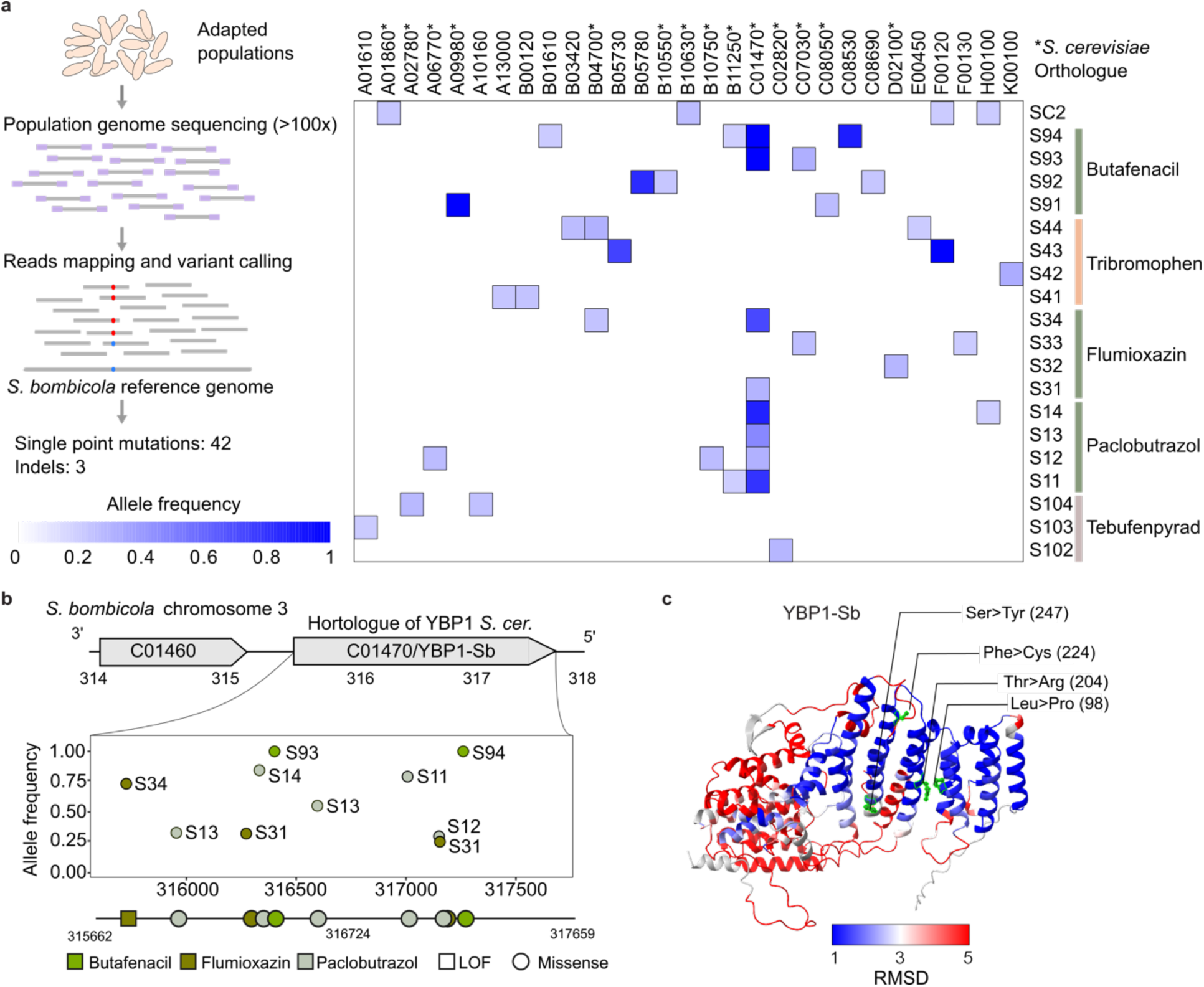
Convergent evolution of *YBP1* during adaptation to pesticides. **a.** Left: Whole genome sequencing strategy used to identify mutations in the evolving lineages. Right: Heatmap showing the *S. bombicola* genes featuring mutation/s in the evolved populations. Colour intensity is proportional to the allele frequency in the corresponding population. Asterisk denotes a gene that has a *Saccharomyces* orthologue. **b.** Genomic region of C01470/YBP1-Sb location showing the detected mutations. For each mutation, the figure shows allele frequency (y-axis), location in kbp (x-axis), chemical stressor (colour), and the type of mutation (shape). **c.** AlphaFold-predicted structure of C01470/YBP1-Sb, coloured by per-residue root-mean-square-deviation (RMSD) relative to the structure of the *Saccharomyces* ortholog, YBP1^Scer^. The ball and stick residues highlight some of the mutations that emerged during the ALE.

One of the populations exposed to Butafenacil gained a mutation in the gene encoding a protoporphyrinogen oxidase (homolog of *YER014W*), falling in its FAD-NADP binding domain. The homolog of this gene is the molecular target of Butafenacil in plants, showing that this compound targets the same enzyme in other species (**Supplementary table 3**) and explaining the ‘off-species’ toxicity uncovered in our screening. The strongest case of convergent evolution is the gene C01470, an ortholog of the *S. cerevisiae* gene *YBP1*, which harboured 10 distinct mutations across 3 different chemicals (Paclobutrazol, Butafenacil, Flumioxazin) and in different populations suggesting a strong selection on the gene function (**Fig 3a right, 3b, Supplementary table 3**). Among the 10 mutations, 9 resulted in missense mutations in amino acids present in alpha helixes of the protein, known to be important in protein-protein interaction (**Fig 3c**). We also detected a frameshift generating a stop-gain at codon 24 of the protein suggesting a loss-of-function as the alpha helixes of the protein are all present after that codon (**Fig 3b-c**). The *Saccharomyces* ortholog of *YBP1* encodes a protein involved in oxidative stress response by interacting with the protein Yap1p. We did not identify an orthologue of *YAP1* in the *S. bombicola* genome, but other *YAP* genes were identified (*YAP5*, *YAP7* and *YAP8*) indicating that interacting proteins are present in the genome. Structural comparison of the two homologues (*YBP1^Scer^* and *YBP1^Sbom^*) based on AlphaFold-predicted structures showed a notable conservation in the N-terminal alpha-helixes of the protein (**Fig 3c**). Structural location of the mutations selected during the adaptive evolution suggests altered function by potential destabilising the alpha-helixes or by partially disrupting the protein-protein interaction interface (**Fig 3c**). Together with the identified stop-gain mutation, this suggests that altered protein interaction upstream the oxidative stress signalling pathway leads to improved chemical resistance.

Together, our ALE experiments and genomic analysis shows that *S. bombicola* can adapt to pesticides and that a common molecular pathway linked to oxidative stress resistance is involved in the susceptibility to different types of pesticides.

### Oxidative stress response and horizontally transferred bacterial genes underpin adaptation to Paclobutrazol

To understand how *YBP1^Sbom^* is mechanistically linked to pesticide resistance, we further evaluated two adapted populations, one evolved in Paclobutrazol and another in Flumioxazin. The former exhibited the highest fitness gain in Paclobutrazol and harboured a missense mutation, while the latter had evolved a loss of function (LOF) mutation. As *YBP1^Sbom^* orthologue in *S. cerevisiae* is linked to oxidative stress resistance^24^, we measured reactive oxygen species (ROS) production when the ancestral and the evolved strains were exposed to Paclobutrazol and H_2_O_2_. For both chemicals, the ROS production was diminished in the evolved strains confirming improved oxidative stress response (**Fig 4a-b, Supplementary table 4**). In accord, mitochondrial biomass was also affected by the oxidative stress. In the ancestral strain exposed to Paclobutrazol, mitochondrial biomass was almost 4-fold higher (t-test, two-sided, p-value=9.57x10^-9^) while the increase in the adapted strains was much lower (1.26-fold for Paclo_evo1, and 1.10 for Flumiox_evo4, t-test, two-sided, p-value=2.14x10^-5^, t-test, two-sided, p-value=6.1x10^-4^) (**Fig 4c, Supplementary table 5, Supplementary Figure 3**).

**Figure 4.**
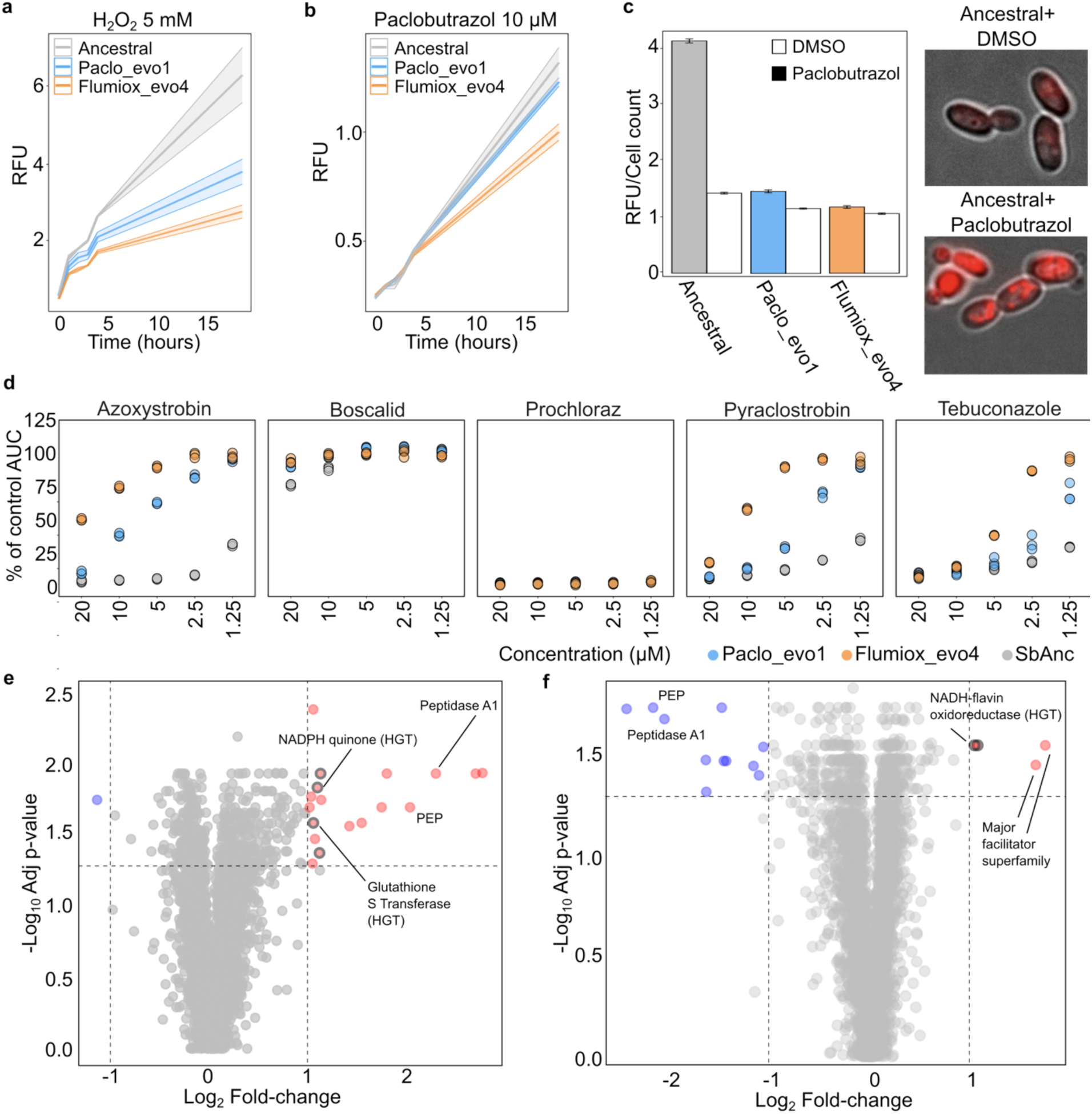
*S. bombicola* response to pesticide is driven by genes involved in oxidative stress response and those horizontally transferred from bacteria. a. ROS production measured as relative fluorescent unit (RFU) of the emission of CM-H_2_DCFDA in ancestral and evolved lines exposed to H_2_O_2_ (5mM). The shaded area represents the standard deviation of each measure at each time-point which is the average for n=4 independent biological replicates. Y-axis reports the RFU measured for each time point. **b.** as in **a.** but for Paclobutrazol exposure. **c.** Quantification of mitochondrial biomass using Mitotracker. Each bar plot represents the average relative fluorescent signal (RFU) of n=4 independent cultures and the error bar the standard deviation. The RFU is normalised to the cell count expressed in optical density. On the right, microscopy images illustrate the cellular response in cultures treated with the Mitotracker (complete images in **Supplementary** figure 3). **d.** Each plot represents the percentage of area under the curve (AUC) compared to the AUC of the control strain grown without chemicals (y-axis). A 100% value means that the AUC, at a given concentration of a compound, is equal to the AUC of the control condition indicating complete resilience to the chemical at that concentration. The ancestral growth is reported for reference of the improvement obtained in the two evolved populations. Each dot represents an individual replicate (n=3). **e.** Summary of proteomic changes comparing the ancestral strain exposed or not exposed to Paclobutrazol. Coloured dots have a significant change of more than 2-fold (ANOVA protein fold change, followed by Benjamini-Hochberg adjusted p-value < 0.05). Grey circles mark genes of bacterial origin that we found to have been horizontally transferred to *S. bombicola* (see main text). Each dot is the average of n=4 biological replicates. **f.** Summary of proteomic changes comparing the ancestral strain with the evolved one exposed to Paclobutrazol. Coloured dots have a significant change (ANOVA protein fold change, followed by Benjamini-Hochberg adjusted p-value < 0.05) of more than 2-fold. Dotted lines in **e** and **f** represent thresholds used for selecting up-regulated and down-regulated genes.

We next tested whether the evolved strains can tolerate other pesticides that are known to act by increasing oxidative stress. We selected 5 additional pesticides not included in our evolution, viz., Azoxystrobin, Pyraclostrobin, Boscalid, Prochloraz and Tebuconazole. All five are found in pollen and honey, a major ecological niche of *S. bombicola*, and are known to act via increased oxidative stress^23^. Consistent with our hypothesis, the evolved strains were cross-protected against four of the five pesticides, viz. Azoxystrobin, Pyraclostrobin, Tebuconazole and Boscalid (**Fig 4d**), attesting the enhanced oxidative stress resistance of the evolved yeasts.

To identify the molecular underpinnings of the enhanced oxidative stress response, we compared proteome-wide changes between the ancestral and the evolved strains upon Paclobutrazol exposure (*n*=4, biological replicates) (**Supplementary table 6**). In the ancestral strain, 18 proteins had increased abundance by over 2-fold (**Fig 4e**), while only one protein had decreased abundance. The upregulated proteins include those involved in countering oxidative stress such as Glutathione S-transferase and NADPH quinone, as well as proteins linked to mitochondria metabolism, including the orthologue of the gene *YNR018W.* Proteins countering proteolytic stress, which can result from oxidative stress, such as peptidases belonging to the A1 family or endopeptidases (**Supplementary Table 7**) also had increased abundance under Paclobutrazol exposure. We further found proteins linked to cell wall stress being up-regulated (**Supplementary Table 7**). When we compared the ancestral strain to the evolved one, both in the presence of Paclobutrazol, the evolved strain featured increased abundance in 2 major facilitator superfamily transporters and 2 other proteins linked to oxidative stress response (**Fig 4f**, **Supplementary Table 8**). The proteins in metabolism and cell-wall stress were not altered in the evolved strain in accord with its increased fitness in the presence of Paclobutrazol and therefore resulted as lower abundance compared to the ancestral strain.

Interestingly, we noted that 4 of the proteins with increased abundance in the ancestral strain and 2 in the adapted strain are of bacterial origin. We first confirmed that these bacterial proteins are not contaminants and the corresponding genes are present also in the reference genome (**Supplementary Figure 4**). Bioinformatic analysis confirmed that these proteins have been horizontally transferred to *S. bombicola*. The respective genes are annotated as having bacterial origins by eggnog, while the other genes are annotated to yeast as expected in accordance with the YGAP pipeline (**Supplementary Table 7,8, methods**). As an additional validation, we performed phylogenetic analysis in accord with the previous work on horizontal gene transfer^25^. We found that the phylogeny of bacterial-origin of the NADPH-quinone oxidase, an important protein in mitochondrial oxidative stress, is consistent with inheritance from bacteria; it branches away from another yeast (*Saccharomyces*, outgroup) and resides within bacterial species that represent the top blast hits whereas no yeast species was among the best 100 hits (**Supplementary Figure 5, methods**).

Together, phenotypic and proteomic data show that mitochondrial oxidative stress is the main driver of paclobutrazol toxicity, and adaptive increase in the abundance of yeast and bacteria-origin proteins confer resistance to oxidative stress caused by several pesticides.

### *S. bombicola* ameliorates negative behavioural changes in bees in response to Paclobutrazol

To investigate the ecological relevance of our laboratory-evolved *S. bombicola* strains, we tested the effects of Paclobutrazol on honeybees when feeding the ancestral and the adapted strains. We monitored two key behavioural phenotypes, sucrose responsiveness and non-associative learning, as these are known to be disrupted by pesticides^26^. Individuals of *Apis mellifera ligustica* were collected from 4 different colonies and randomly assigned to either the treated (pesticides and/or yeast) or the untreated controls (DMSO and/or glycerol) (**see methods**).

We first assessed the toxicity of paclobutrazol to honeybees at 3 different concentrations: 25, 130 and 340 µM (**Fig 5a, top**). The two higher concentrations represent the commercial pesticide formulation. Further, pesticides can accumulate in provisions during foraging depending on the lipophilicity expressed as logarithmic octanol water coefficient (logK_ow_). Compounds with logK_ow_ between 2.5 and 4.7 are reported to accumulate in beeswax, and Paclobutrazol have a logK_ow_ of 3.11 according to the Pesticides Properties Database (PPDB). In addition, pesticides can get concentrated in the natural environment because of evaporation. Paclobutrazol is classified as persistent and its reported half-time degradation DT_50_ can vary between 27-61 days up to 13 months according to the PPDB. Therefore, concentrations can remain high or increase even without direct exposure because of bioaccumulation in the hives or as a natural persistence. We observed that the two high doses significantly shifted the sucrose habituation curves reflecting increased appetitive motivation, which is indicative of the stress triggered metabolic change and increased energy requirements (non-associative learning test, **Fig 5b**, GLMM test p-value<0.0001, Post-hoc Dunnett test, Control-High: p=0.007, Control-Medium: p=0.01). This is also reflected in the habituation score (Habituation score (HS), Kruskal-Wallis p-value<0.001, **Supplementary Tables 9 and 10**). Similarly, the sucrose response score (SRS), which indicates the lowest sugar concentrations at which a bee still feeds, increased upon pesticide treatment (Kruskal-Wallis p-value<0.001, Post-hoc Dunnett test, Control-High: p=0.013, Control-Medium: p=0.034, Control-Low: p>0.05), reflecting enhanced bee responsiveness even at low sugar concentrations. This alters the colony’s division of labour and reduces overall foraging efficiency^27^ (**Supplementary Tables 9 and 10**). These responses have been observed for other pesticides^26^ and are a hallmark of the pesticide toxicity as the bee needs more energy to counteract the toxic effect ^28^.

**Figure 5.**
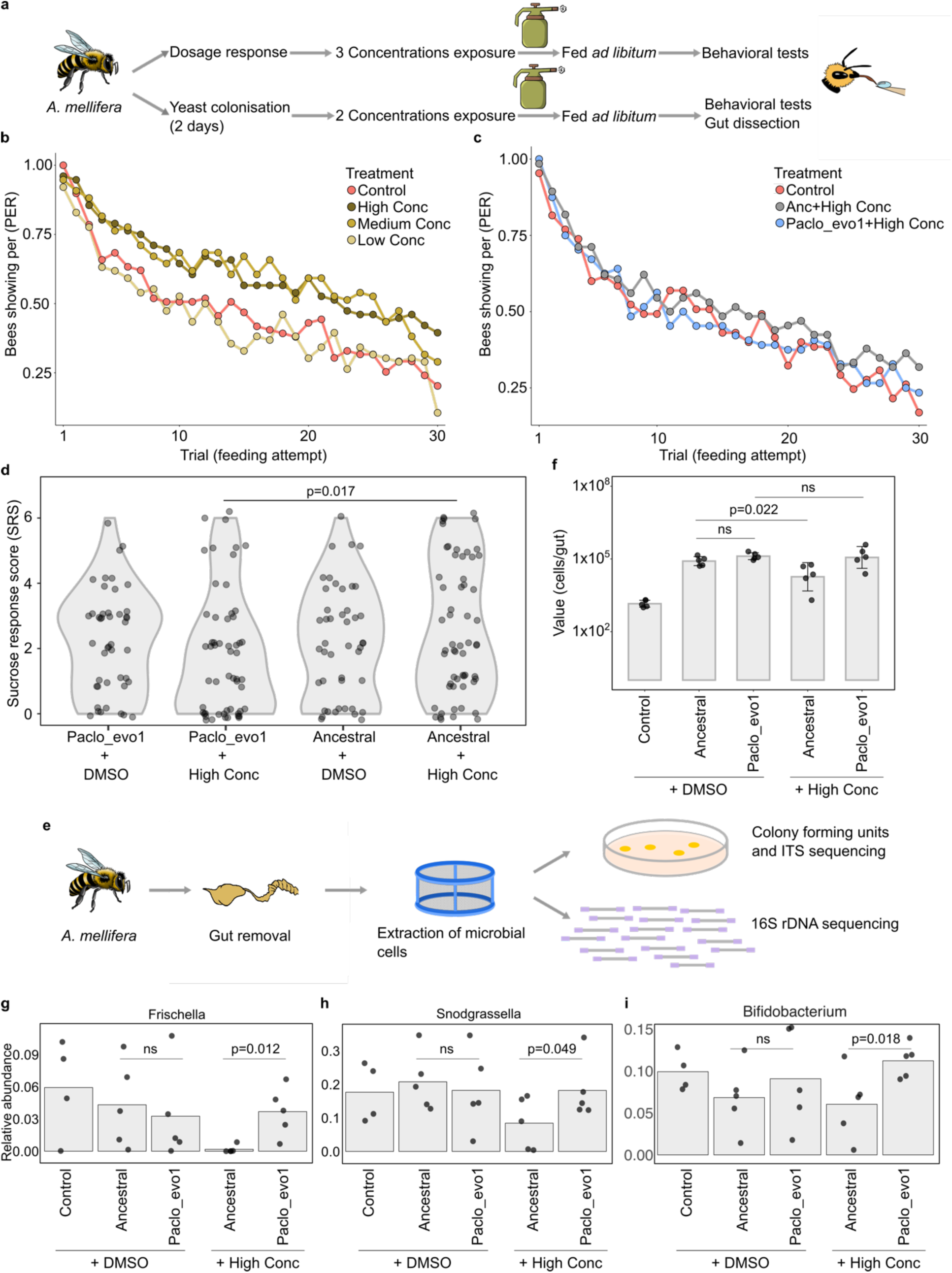
Yeast colonisation modulates pesticide impact and gut microbial abundance in honeybees. **a.** Experimental design of the *in-vivo* experiments. **b.** Habituation curve to feeding stimuli (x-axis) of honeybees exposed to the three dosages of pesticides (low=25 µM, medium=130 µM, high=340 µM). The y-axis reports the ratio of the bees in each treatment group which performed the proboscis extension reflex (PER) upon stimulation. (n=76 for low, medium, and high dose; n=79 for the control) **c.** Habituation curve to feeding stimuli (x-axis) of honeybees exposed to the 340 µM Paclobutrazol to which were fed the Ancestral (Anc) or the evolved (Paclo_evo1) yeast. The y-axis reports the ratio of the bees in each treatment group which performed the proboscis extension reflex (PER) upon stimulation (Control n=74, Paclo_evo1+high dose n=74, Anc+high dose n=72). **d.** Sucrose response score (SRS) as the number of concentrations of sucrose to which the honeybees responded highlighted the difference between the bees exposed to the high concentration of pesticide **e.** Schematic overview of the gut dissection and extraction of microbial cells through dedicated filters. Cells were plated on selective growth media to estimate the yeast total abundance. Single colonies were profiled using ITS to confirm species-level identity for yeasts (top). Bacterial abundance was estimated using 16S sequencing **f.** Results of the colony forming unit assay (CFU) on selective media plating as cells/gut (y-axis, log_10_ scale) to evaluate yeast population size in the honeybees fed with the yeasts and with or without the pesticide exposure (n=5 biological replicates). The error bars represent the standard deviation while each dot is an individual dissected gut. **g.** Bar-plot showing relative abundance of *Frischella* across the samples (n=4 for Control, n=5 for other treatments). Each dot represents a single gut sample, while the bar height represents the average of the abundance across the samples. (t-test one-sided p-value=0.012, Ancestral versus Evolved + High Conc) **h.** Bar-plot showing the relative abundance of *Snodgrassella* across the treated samples. Each dot represents a single gut sample, while the bar height represents the average of the abundance across the samples (t-test one-sided p-value=0.049, Ancestral versus Evolved + High Conc). **i.** Bar-plot showing relative abundance of *Bifidobacterium* across the treated samples. Each dot represents a single gut sample, while the bar height represents the average of the abundance across the samples (t-test one-sided p-value=0.018, Ancestral versus Evolved + High Conc).

To test whether *S. bombicola* could rescue the pesticide-induced behavioural changes, we fed the evolved and ancestral populations to the honeybees ad libitum for 2 days before exposure to Paclobutrazol (**Fig 5a**, **see Methods**). The yeast feeding did not alter honeybee survival compared to the control confirming that this yeast does not negatively affect its host at this concentration (**Supplementary Figure 6a**). We then performed the two behavioural tests following exposure to two commercial formulation equivalent doses of pesticides (**Supplementary Table 11 and 12**). Yeast feeding mitigated the pesticide-induced behavioural alterations. Both the evolved and the ancestral strains restored the proboscis extension reflex (PER) in the habituation test to levels comparable to the control group at the high pesticide treatment (GLMM p-value>0.05, **Fig 5c**) and the habituation scores (HS) (Kruskal-Wallis test p-value=0.55). We also observed an improvement in the sucrose responsiveness test for both yeasts (SR, based on the percent of bees with PER), and a marginally significant difference between the ancestral and evolved yeast fed bees (GLMM p-value=0.054) (**Supplementary Figure 6b**). Notably, bees fed with the evolved yeast cells showed a significantly lower sucrose response score (SRS) than those fed with the ancestral (Ancestral versus Paclo_evo1, p-value=0.017, t-test one sided) (**Fig 5d**). These behavioural tests show that yeast feeding can ameliorate behavioural defects caused by Paclobutrazol.

### *S. bombicola* colonises the honeybee gut and modulates bacterial composition

To confirm that the positive impact of *S. bombicola* feeding was due to colonisation of the gut and not by a nutritional value conferred by the yeast, we dissected the honeybee guts and plated its content on selective growth medium to allow growth of only yeast colonies (**methods**). Indeed, the total yeast population density was ∼100 times higher in bees fed with *S. bombicola* than in the control indicating that the cells actively colonised the gut (**Fig 5e-f, Supplementary Table 13**). To test whether the colonies were that of *S. bombicola*, we sequenced the ITS genomic regions of randomly selected isolates and found that around 90% (22 of 25 tested) of the colonies were *S. bombicola*. In control, non-exposure, experiments, bees fed with evolved strains showed similar colonisation levels to that of the parental lineage (t-test, two-sided p-value=0.67) (**Fig 5e**). In contrast, under Paclobutrazol exposure, the population of ancestral strain significantly dropped (t-test, two-sided p-value=0.022, 3∼fold average drop) while that of the evolved strain remained stable (t-test, two-sided p-value>0.05) (**Fig 5f**). Thus, increased colonisation under Paclobutrazol exposure attested that the laboratory-evolved strain maintains its pesticide resistance *in vivo* and outperformed the ancestral strain under chemical stress.

Pesticides treatment also caused a major impact on the total microbial abundance (**Supplementary Figure 6c-d**). To assess how yeast feeding and Paclobutrazol exposure affected bacterial population in the gut, we used 16S rRNA amplicon sequencing (**Fig 5e**). We could identify all the known major genera of the bee-gut microbiome in our samples indicating that the microbiomes of the bees used are comparable to those in previous studies^29,30,31^ (**Supplementary Figure 8a, Supplementary Table 16**). Qualitatively, feeding with ancestral or evolved yeasts did not show any difference in the community membership (species presence), except for some differences in very low abundance species (Jaccard-distance p-value<0.05, **Supplementary Figure 7**). We then evaluated the variability in the relative abundance of bacterial species upon the pesticide exposure when feeding the ancestral or the evolved yeast. The bees fed with the laboratory-evolved strain had a higher abundance of beneficial bacteria, *Frischella* and *Snodgrasella* to those fed with the ancestral yeast (**Fig 5g-h**). No such difference was observed in the absence of pesticide treatment suggesting that the boost from the evolved yeast is linked to its pesticide resistance. The bacteria from *Frischella* genus have been reported to feature positive effect on hosts immunity, while *Snodgrasella* is involved in pathogen resistance and cross-feeding metabolites between host and the other members of the microbiota^30,32^. Feeding with the evolved yeast also led to higher abundance of *Bifidobacterium* (**Fig 5i**) which are generally considered beneficial^30^. Together, these results indicate that the evolved yeast can stably colonize the honeybee gut under chemical stress and its beneficial effects extend to the bacterial members of the bee gut microbiota.

## Discussion

Our results show that many non-fungicide chemicals, such as insecticides and herbicides, impact the growth of various yeast species, spanning the Saccharomycotina sub-phylum, which play an important role across many ecosystems, from pollinators to human health^33,11^. Published eco-toxicological data indicates ubiquitous presence of many of these chemicals, including that in rivers, pollen, and nectar or hives. In case of some chemicals, including Paclobutrazol, the reported environmental concentrations are higher than those used in our evolution experiments^23,34^. Further, a recent study suggests that environmental contamination for many compounds is more widespread than currently documented due to limited analyses and analytical considerations^35^. Our systematic screening of chemical-yeast interactions thus identifies yeasts as an off-target of common pesticides, many of which are non-fungicides.

Amongst the diverse yeast species that we tested, *Starmerella bombicola* stood out with broad sensitivity to many chemical pollutants. Association of this species with many bees^17,15,18^ raises the question of impact on critical pollinators through their microbiomes. Answering this question requires understanding the molecular mechanisms underlying the chemical sensitivity. Laboratory evolution under controlled conditions helped us to identify regulation of oxidative stress as the key determinant common to many chemicals. The orthologue of *YBP1* emerged as a convergent evolutionary target of pesticides adaptation. The orthologs of this protein in other yeast species interact with the YAP proteins to regulate resistance to diverse stresses^36^.

We also discovered bacterial genes obtained through horizontal gene transfer (HGT) in *S. bombicola* that play a role in the resistance to pesticides. Their role in oxidative stress resistance suggests that these HGT genes are highly integrated into the physiological network of *Starmerella*. Some of these genes are present in other *Starmerella* species and therefore it is unlikely that the acquisition of these bacterial genes is a recent adaptation to pesticide use. The acquisition is likely linked to the ecological niche occupied by this yeast, as the family of bacteria donors are also present in the bee microbiome. One ecological explanation is that the acquisition of these genes from bacteria has been selected for countering oxidative stress, e.g., when moving between the host and non-host environments.

Could the beneficial adaptations emerging in our laboratory experiments occur in nature? While *S. bombicola* was able to adapt to many chemicals, our experiments were carried out under well controlled conditions. In natural settings, adaptation to specific chemicals may be hampered due to multiple selections acting simultaneously. Moreover, evolution in nature may be constrained by the population size, which in the bee gut (10^1^ -10^4^ rarely up to 10^7^) and flowers (10^1^-10^5^) is lower than those of our experimental evolution^19,37^.

Oxidative stress is a common consequence of pesticide exposure in honeybees^38,39^ and other bees^40^. Our data shows that the bee microbiome members, especially like *S. bombicola*, contribute to the countering of this stress and laboratory evolution can further boost this beneficial effect. Yeast supplementation was able to modulate deleterious pesticides effect on bee behaviour, which is commonly altered by pesticides exposure^26,40,41^. The beneficial effects were also evident in maintaining bacterial genera linked to positive host interactions. How the yeast promotes the maintenance of beneficial bacterial genera need further research, but other studies have highlighted a positive association between yeast and *Gilliamella* and *Snodgrassella* in the honeybee gut^19^ or more in general between fungi and bacteria^42^, or a beneficial effect of yeast derived metabolites on the host^43^.

Our results open a possibility for leveraging natural bee-associated microbes for mitigating harmful effects of pesticides. While microbiome interventions are considered promising for addressing anthropogenic impact on ecosystems, their implementation is hindered by lack of mechanistic insights and regulatory and ethical considerations^44^. Our study demonstrates how laboratory evolution under controlled conditions can overcome these hurdles: evolution promoted specific resistance in natural isolates without compromising colonisation fitness and with a detailed understanding of the mechanisms involved. Further, the evolved yeasts are non- genetically modified organisms easing the regulatory path for their deployment in the environment. Taken together, our results open a mechanistic window into chemical-yeast-host interactions and bring forward laboratory evolution as an effective approach to mitigate effects of pesticides on pollinators.

## Materials and Methods

### Yeast strains and growth media

Most yeast strains were obtained from the UK’s National Collection of Yeast Cultures (NCYC) with accession codes listed in **Supplementary Table 1**. Other strains were previously isolated in our laboratory from Kefir^45^. All cultures were grown in/on YPD (yeast extract peptone dextrose media) with a final composition of 10 g/L yeast extract (BD Bacto), 20 g/L peptone (BD Bacto), 20 g/L glucose (Sigma-Aldrich, Cat# G8270) and optionally 20 g/L agar (BD Bacto) in polished/milliQ water. Glucose was prepared as a 40% stock and autoclaved separately from the other components. YPD ensure that all yeasts used in this study spanning different phylogenetic groups can grow as reported from the NCYC guidelines. All experiments were conducted at 30 °C unless specifically defined.

### Growth screening

Assay plates were prepared by diluting the pesticide library aliquots (2 mM in DMSO) 50-fold in YPD within a deep-well plate from which 100 μl were aliquoted into 96 square-well plates (Enzyscreen). Liquid handling was performed using a Biomek i7 Automated Workstation (Beckman Coulter).

Cells were revived from glycerol-preserved cryostocks on YPD agar plates, which were incubated for 1-4 days. From these, 4 ml liquid cultures were inoculated and grown overnight with shaking (150 rpm). The optical density at 600 nm (OD_600_) was determined and cultures were diluted in 20 ml of fresh media to a final OD_600_ of 0.1. After approximately 6 h, cultures were again diluted to OD_600_ 0.1 and 100 µl were added to each well of the previously prepared assay plates (resulting in a starting OD_600_ of 0.05 and a compound concentration of 20 μM). Assay plates were sealed with breathable, opaque seals (Merck, Z763624-100EA). Three biological replicates were grown and measured on separate days. Growth curves were recorded using a GrowthProfiler instrument (Enzyscreen), which records photographic images of the plates through their transparent bottom, from which the turbidity of the culture can be estimated. The following settings were used: temperature - 30°C; shutter time - 4 ms; interval between measurements -60 min; shaking speed - 150 rpm. The AUC reduction at different concentrations was performed following the same protocol.

### Growth screen data analysis

A command line version of the Growth Profiler Viewer (Enzyscreen, v. 2022_03_03) was used to crop individual wells from each image and extract the green value which served as biomass proxy. The following settings were used: well distance in x-direction - 9 mm; well distance in y-direction - 9 mm; well shape - square; spot shape - rectangle; size spot in x-direction - 3 mm; size spot in y- direction - 3 mm; deviation from centre in both directions - 0 mm; geometric correction - 0.5 mm; ignore top G pixel - 50%; ignore bottom pixel - 10%; compensate for shutter time - no. Values were exported and further analysed in Python. Growth curves were baseline-corrected by subtracting the initial value from all measurements. Growth curve parameters were then extracted with pyphe^46^, using the pyphe-growthcurves command line tool as described in ^47^ with the following settings: lag-threshold - 1.25; t0-fitrange - 1. Normalisation and data aggregation was performed with pyphe-analyse, using ‘sum of values (AUC)’ as the growth curve parameter to estimate fitness. AUC values were normalised to the plate median, resulting in relative values where 1 indicates no compound effect.

Hits were called for every replicate separately by subtracting the plate-corrected AUC of the vehicle controls from that of the compounds, followed by division of the difference by the standard deviation of the controls. This z-score like value was converted to p-values using the survival function implemented in scipy.stats.norm.sf and the result was multiplied by 2 for two-sided testing, followed by FDR correction for testing of multiple compounds using the Bonferroni method (performed on each species and replicate separately). An additional threshold of at least 15% reduction of growth relative to the controls (plate-corrected AUC < 0.85) was applied to filter out statistically significant, yet small and biologically less meaningful reductions in growth. A compound was considered a hit globally, if it was a hit (p_adj_ <0.05) in at least two out of the three biological replicates. One compound was excluded from the original list of 1025 compounds for a contamination of the stock in the library and therefore was excluded for further analysis.

### Adaptive laboratory evolution (ALE) of *Starmerella bombicola*

For the ALE experiment 4 independent populations of *S. bombicola* (NCYC 2908) were inoculated from a pre-culture in YPD into wells of a 2 mL deep-well bottom plate (ThermoScientific Cat# AB0661) at a starting OD_600_ of 0.05 each containing a total volume of 1 mL of YPD for a total of 44 independent populations consisting of 4 replicates for each of the individual ALE conditions including the DMSO control. Information on the pesticides used was retrieved by the Pesticides Property Database https://sitem.herts.ac.uk/aeru/ppdb/. The strains were grown for 48 hours until and then the OD was measured to evaluate population size at each passage until passage 18 and used to calculate the transfer inoculum to adjust it to be around OD_600_ 0.05 while the last transfer was only used to prepare the cyrostock. For assessing improvement, the OD data were normalised to YPD controls to take into account experimental variability in OD measures. The whole process was replicated for 20 transfers and each 5 transfers a frozen cryostock of the culture was stored at -80 °C by adding glycerol to a final concentration of 25% directly in the plate used for the transfer.

### Genome sequencing analysis of evolved populations and gene annotations

DNA extraction from the evolved populations was performed using the kit DNA ZymoBIOMICS miniprep kit (Zymoresearch™, Cat# D4300) following manufacturer protocol. DNA was sent for sequencing at the Genomics core facility at EMBL, Germany and 150-bp pair reads were generated on a Novaseq2000 using P2 chemistry. The fasta files were analysed for quality control using FastQC (v0.12.0). Following, the Illumina reads were mapped against the reference genome for NCYC2908 downloaded from the NCBI database (Ref: GCA_001599315.1) and indexed using bwa (v0.7.18) with the flag ‘-mem’. Mapped reads were then processed using samtools (v1.9) ‘markdup’ option to remove optical duplicates and the depth of coverage was calculated using ‘samtools depth’. Variants against the reference genome were called using Freebayes (v1.3.2) using a threshold of --min-coverage 30 and selecting variants with an allele frequency of at least 20 % to select adaptive variants emerging in the populations with a considerable frequency and using the flags ‘-C 10’ and ‘--pooled-continuous -p100’. Vcf files were then processed using bgzip and tabix and common variants between all control samples and independent evolved populations were filtered out using bcftools (v1.9) ‘isec -C’. Filtered Vcfs were then pooled together, and non-unique variants were removed as mutations occurring in the same position with high frequency and same genotype and likely false-positive due to pre-existing variants. Annotation of the genomic region was performed using an ad-hoc R script that maps the position of the variant to the annotated genomic region identified with the YGAP pipeline^48^.The YGAP pipeline utilises for gene annotation a synteny based approach mapping to other *Saccharomycetacae*. Genes without orthology information are also annotated but there is no reference orthologous gene name based on *Saccharomyces* yeast gene nomenclature. Genes were named using the nomenclature relative to the *S. bombicola* genome and were useful for interpretation the *Saccharomyces* orthologue was reported. For the annotation of the effect of the identified variants by Freebayes, SnpEff (v5.2-1) was used on the generated vcfs. IGV (v2.19.7) was used to inspect variants identified by Freebayes in the *YBP1* gene.

### AlphaFold structure construction

Protein structures were constructed using AlphaFold3^49^ for Ybp1p from *Saccharomyces cerevisiae* (strain ATCC 204508 / S288C) and its homolog SBOM0C01470 from the reference genome of the strain used for the ALE experiment, as no experimentally determined structures were available. Structural alignment was performed using the “matchmaker” tool in ChimeraX v1.9^50^, based on root-mean-square deviation (RMSD) values with the default settings. Residue mutations were visualized as ball-and-stick models, while the rest of the protein structure was rendered in cartoon representation.

### ROS analysis, Mitochondrial content estimation and microscope imaging

For measuring the ROS production of the ancestral and evolved lines, cells were first grown overnight in 5 mL YPD at 30 °C shaking at 220 rpm. The following day cells were resuspended at an OD_600_of 0.8. Cells were let grown for 4 hours at 30 °C with shaking at 220. For measuring ROS production H_2_O_2_ was added at 5 mM as positive control and fluorescence of the dye CM-H_2_DCFDA at 10 μM (ThermoFisher, Cat# C6827) was measured using a VarioskanLUX (ThermoFisher) plate reader at 0, 1, 2, 3, 4 and 19 hours in the H_2_O_2_ treated and Paclobutrazol treated samples. For estimating mitochondrial content, the dye MitoTracker™ Green FM (ThermoFisher) was used at a final concentration of 200 nM and added to the cells treated for 4 hours with Paclobutrazol 30 minutes before the fluorescence measure, while for microscopy images MitoTracker™ Red FM was used at the same concentration. The fluorescence of the cells stained with the MitoTracker was measured using a VarioskanLUX plate reader with excitation/emission wavelength of 490/516 nm. An aliquot of the treated cells with MitoTracker was used to estimate population size with a benchtop spectrophotometer to normalise the fluorescence readout to the number of cells in the culture.

For microscope imaging yeast suspensions were diluted to an OD_600_ of 0.8, stained with 200 nM MitoTracker™ Red FM for 15 minutes, then centrifuged and washed with PBS before imaging. Images were acquired using a transmission light microscope (Zeiss Axio Imager; Carl Zeiss, Germany) equipped with a 16-bit CMOS camera (Hamamatsu ORCA-Flash 4.0, 6.5 µm/pixel) and controlled with Zen 2011 software. Single images were collected with a 100× oil-immersion objective (Plan-Neofluar 100×/1.30; Zeiss), yielding a field of 2048 × 2048 pixels at 0.065 µm/pixel. Phase contrast and fluorescence modes (excitation 579 nm, emission 599 nm) were used with a 50 ms exposure time.

### Phylogenetic and genomic analysis of bacterial HGT origin

The sequence of the putative NADPH oxidase with bacterial origin was obtained from the *S. bombicola* annotated genome. The regions coordinates were obtained from the “.gff” annotation file and used to extract the region where to inspect the depth of coverage profile. The regions were manually inspected with IGV (v2.19.7) to confirm that reads at the border of the gene were mapping correctly and there were not artifacts such as mis mapping read pairs indicative of a misplacing of the bacterial gene in the yeast genome. Then, the amino acid sequence was blasted on NCBI using the “blastp” function and the top 100 hits for sequence similarity were downloaded. The sequences were then aligned using MUSCLE with default options and that was used in MEGA (v12) to build a phylogenetic tree using the Maximum Likelihood method and the Jones-Taylor-Thornton model for substitution method. The sequence of the *Saccharomyces* orthologue protein was included as an outgroup.

### Proteomic cultures preparation

Cryostocks of the ancestral and evolved population were inoculated in 5 mL YPD and grown overnight at 30 °C shaking at 220 rpm. The following days cells were transferred in 1mL fresh YPD in 2.2 mL deep well plate as the one used in the ALE at a dilution of OD_600_ 0.05 and grown at 30 °C shaking at 220 rpm in the presence of 1.25 μM Paclobutrazol or DMSO as control. Cells were grown to an approximately OD_600_ of 1 and then were centrifuged at 4000 rpm at 4 °C for 10 minutes. The media was discarded, and the pellet stored at -80 °C for protein extraction.

### Protein Extraction and Digestion

For downstream mass spectrometry analysis, protein samples were processed using the S-Trap sample preparation protocol. A maximum of 100 µg of protein was transferred into a 2-mL microcentrifuge tube, to which 10% SDS was added to reach a final SDS concentration of 4%. Subsequently, 1 M TEAB buffer (pH 8) was added to achieve a final TEAB concentration of 50 mM. If necessary, HPLC-grade water was added to bring the total volume to at least 50 µL. The protein sample was reduced with 250 mM DTT (final concentration: 20 mM) at 50°C for 10 minutes, followed by a 10-minute equilibration at room temperature. Alkylation was then performed using 250 mM IAA (final concentration: 40 mM) for 30 minutes at room temperature in the dark.

The sample was acidified by adding 12% phosphoric acid to reach a final concentration of 1.2%. Protein precipitation was induced by adding seven volumes of S-Trap binding buffer (90% methanol in 100 mM TEAB, pH 7.1). The solution was gently inverted or vortexed to mix, and visible protein colloid formation confirmed efficient precipitation. Depending on the protein quantity, either S-Trap micro spin columns (≤100 µg) or mini spin columns (100–300 µg) were used, each placed into a 2.0-mL flow-through collection tube. The acidified lysate/binding buffer mix was added (100 µL for micro columns or 200 µL for mini columns), and the columns were centrifuged at 4,000 x g for 10 seconds until the solution fully passed through. The flow-through was discarded and the step was repeated until the full volume was loaded.

The columns were washed twice with 200 µL (micro) or 400 µL (mini) of S-Trap buffer, centrifuging at 4,000 x g for 10 seconds each time. After a final spin to remove the residual buffer, the columns were ready for digestion. Sequencing-grade trypsin was prepared at a 1:25 (w/w) enzyme-to-protein ratio in 50 mM TEAB. The columns were spun to dryness and transferred to clean low-retention elution tubes. Digestion was initiated by adding 125 µL of the trypsin solution to each column. The micro columns were loosely capped and mini columns sealed, followed by incubation at 47°C for 1 hour. A second aliquot of trypsin solution was added, and digestion proceeded overnight at 37°C without agitation.

Peptides were sequentially eluted into a single 2.0-mL tube using three elution buffers to ensure comprehensive peptide recovery. The first elution used 80 µL of 50 mM TEAB (pH 8), the second 80 µL of 0.5% formic acid, and the third 80 µL of 50% acetonitrile with 0.5% formic acid. Each elution step involved centrifugation (1,000 x g for the first two, 4,000 x g for the third). All eluates were pooled, vacuum-dried, and stored at –80°C until further processing.

### TMT Labeling and Peptide Fractionation

The dried peptides were reconstituted in 0.2% formic acid, thoroughly mixed, and desalted using C18 solid-phase extraction columns. After desalting, peptides were dried, quantified, and resuspended in 100 mM TEAB (pH 8.5) to a concentration of 50–100 µg. TMTpro 16plex reagents (Thermo Fisher Scientific) were dissolved in 41 µL anhydrous acetonitrile and added to the corresponding peptide samples at a 1:1 (w/w) ratio. Labeling was performed for 1 hour at room temperature and quenched with 8 µL of 5% hydroxylamine for 15 minutes. Labeled peptides were pooled, dried, reconstituted in 0.1% formic acid, desalted, and dried again.

Offline fractionation was performed using the Pierce™ High pH Reversed-Phase Peptide Fractionation Kit. Labeled peptides were reconstituted in the kit’s wash solution and loaded onto spin columns. Elution was performed stepwise using increasing concentrations of acetonitrile in triethylamine-containing buffer. Eight fractions were collected, dried, and stored at –80°C.

### LC-MS/MS Analysis and Data Processing

Peptides were analyzed on a UltiMate™ 3000 LC system coupled to an Orbitrap Eclipse mass spectrometer operating in data-dependent acquisition (DDA) mode. Peptides (750 ng) were trapped on a PepMap™ Neo C18 trap column and separated on an Acclaim PepMap 100 analytical column (75 µm × 50 cm) at 300 nL/min with a 180-minute gradient: 2% B to 40% over 120 min, 40% to 90% over 45 min, to 95% in 0.1 min, then down to 5% over 15 min. Mobile phases included 0.1% formic acid in water (A) and 80% acetonitrile with 0.1% formic acid (B).

Full MS scans were acquired at 120,000 resolution (m/z 370–1500, AGC target: 300%, max injection time: 50 ms), followed by MS/MS at 30,000 resolution (AGC: 100%, max injection time: 45 ms, NCE: 26). Precursors with charge states 2–5 and intensity ≥ 5.0 x 10³ were selected with a dynamic exclusion time of 65 seconds.

Post-translational modifications were identified using Proteome Discoverer v3.1 with SEQUEST HT, searching against the in-house yeast database proteome (the one annotated with YGAP pipeline mentioned in “Genome sequencing and analysis”). Variable modifications included methionine oxidation, protein N-terminal acetylation, and TMT labeling. A target-decoy approach with a 1% false discovery rate (FDR) was used for PSM, peptide, and protein validation. Identified proteins were then analysed on R and proteins with abundance change higher than 2-fold in each direction and with a p-value <0.05 were considered as significant hits. These sets of proteins were then reannotated using eggNOG-mapper v2 for identifying possible genes horizontally transferred from bacteria to this yeast as the YGAP pipeline annotated some of the identified genes as “hypothetical protein” but not found an ortholog in other yeasts.

### General procedure and bee exposure

To evaluate the effects of the pesticide Paclobutrazol on the behaviour of honeybees (*Apis mellifera ligustica*), foraging individuals were collected as they exited the hive from an apiary located at the Department of Biology, University of Florence. Bees were taken from four different colonies. Immediately after collection, bees were transported to the laboratory, cold-anesthetized for approximately 5 min, and individually harnessed in plastic tubes^51^. A small strip of duct tape was placed between the head and thorax, and the head was secured to the holder using a drop of low-melting-point wax, allowing free movement of the antennae and mouthparts only. After a recovery period of about 30 min, bees were randomly assigned to either control or treatment groups and fed accordingly. Control bees received 5 µl of a 40% sucrose solution mixed with DMSO at a 10:1 volume ratio. Treated bees received 5 µl of the same sucrose-DMSO solution containing Paclobutrazol at one of three concentrations: High (H: 340 µM), Medium (M: 130 µM), or Low (L: 25 µM). Following treatment, all bees were fed *ad libitum* with a 40% sucrose solution to standardize hunger levels and were kept overnight in a dark, humid environment at room temperature (23 ± 2 °C)^51^. The proboscis extension reflex (PER) can be elicited in restrained bees by stimulating the antennae with a sucrose solution. Bees were tested the following day according to the respective experimental protocol (see below).

For the yeast experiment (Y1=Paclo_evo1, Y2=Ancestral), bees were collected as previously described and immediately housed in groups of 50 individuals within small plexiglass cages (9 × 7 × 11 cm), maintained at room temperature (24 ± 2°C) and relative humidity of 50 ± 10%. Each cage was fitted with a 20 ml syringe to provide either the experimental or control solution *ad libitum* for two days. Three treatment groups were established: one group received Y1 (in suspended in 2.5% glycerol in 40% sucrose solution), a second group received Y2 (suspended in 2.5% glycerol in 40% sucrose solution), and a control group was provided with 40% sucrose solution supplemented with 2.5% glycerol only. Cell concentration for the colonisation was similar as ^16^ and in the range of 5.5x10^6^ cell/mL. Bee survival was recorded daily during the two-day exposure period. On the third day, bees from the Y1, Y2, and control groups were individually harnessed in plastic tubes as described above. Each bee was fed 5 µl of 40% sucrose solution with DMSO (10:1 vol/vol; control) or with DMSO plus Paclobutrazol at one of two concentrations: High (H: 340 µM) or Medium (M: 130 µM). Following treatment, bees were given *ad libitum* access to 40% sucrose solution to standardize hunger levels and were kept overnight in a dark, humid environment at room temperature (23 ± 2°C). Bees were tested the following day according to the appropriate experimental conditions (see below).

### Sucrose responsiveness

Prior to conducting the sucrose responsiveness assay, bees were repeatedly stimulated with water and allowed to drink as needed, thereby reducing the likelihood of eliciting a PER to water rather than to the sucrose solution itself. An additional water stimulus was administered at the start of the assay, and any bees that responded were excluded from further testing^27,52^. Sucrose responsiveness was assessed by sequentially presenting six sucrose solutions of increasing concentrations: 0.1%, 0.3%, 1%, 3%, 10%, and 30%, with a 2-minutes inter-stimulus interval^52,53^. Each sucrose presentation was followed by a water stimulus to prevent sensitization to sucrose. To account for possible lateral differences in sucrose response, both antennae were stimulated with a toothpick in each trial^53^. Bees that failed to respond to any of the test concentrations were also presented with a 50% sucrose solution. Individuals that did not respond were excluded from the analysis, as non-responsiveness at such a high sucrose level may indicate a motor impairment rather than a lack of gustatory sensitivity ^52^. The number of bees excluded from the analysis did not differ significantly between groups (χ^2^ test p-value=0.2) (Control n=11, Y1+high concentration n=14, Y2H+high concentration n=7). PER responses were recorded for each sucrose trial. An individual Sucrose Responsiveness Score (SRS) was calculated as the total number of PERs observed across the six sucrose concentrations, yielding a score between 0 and 6.

### Habituation assay

During the training phase, restrained bees underwent 30 successive antennal stimulations with a 10% sucrose solution, each lasting less than one second and separated by a 10-s inter-trial interval^51^. The presence or absence of the PER was recorded for each stimulation. An individual Habituation Score (HS) was then computed for each bee, corresponding to the total number of PERs exhibited across the 30 trials. Bees that failed to respond to the initial stimulation were excluded from analysis, resulting in possible HS values ranging from 1 to 30. The number of bees excluded from the analysis (HS=0) did not differ significantly between groups (χ^2^ test p-value=0.1) (Control n=9, Y1+high concentration n=9, Y2H+high concentration n=6). To assess dishabituation, a single antennal stimulation using a 50% sucrose solution was administered 10 seconds after the final habituation trial (dishabituation trial, DT). Then, after another 10-s interval, bees were re-exposed to the original training stimulus (10% sucrose solution) to determine whether the PER re-emerged. This procedure allowed us to distinguish genuine habituation from other factors such as sensory fatigue or sensory adaptation^54^.

### Statistical analysis of bee experiments

Statistical analyses were performed in R (version 4.4.3). Bee survival during the yeast-only exposure was evaluated using a standard Cox proportional hazards regression model (survival package). A Chi-squared tests evaluated differences in the proportion of excluded individuals (those failing to respond to 50% sucrose or scoring zero in habituation). For all experiments involving pesticide exposure and combined yeast and pesticide exposure, sucrose responsiveness and habituation were analysed using generalized linear mixed-effects models (GLMMs; binomial distribution, logit link) via the lme4 package. Models included treatment and either sucrose concentration or trial number as fixed factors, and bee identity nested within colony (Colony:ID) as a random effect. Post-hoc comparisons against the control were performed using Dunnett’s contrasts via the glht function (multcomp package). Overall Sucrose Responsiveness Scores (SRS) and Habituation Scores (HS) were compared using Kruskal-Wallis tests, followed by Dunn’s post-hoc tests with Holm’s p-value adjustment for pairwise comparisons. A paired *t*-test was used to specifically compare SRS between the ancestral (Anc) and evolved (Paclo_evo1) yeast groups.

### Guts Dissection and microbial population counting

Guts stored in 25% glycerol were let thaw in ice then each gut was transferred to a 40 µm pore size filter (Cat# 732-2757) and forced through the mesh of the filter with a sterile syringe plunger gently to dissolve the gut and release the microbial cells into a well of a 6-well plate on which the filter was fitted. The filter was washed twice with 500 µL of the same solution of glycerol in which the gut was stored to collect the cells in a cell suspension of a total volume of 1 mL. Then 20 µL of the solution were serially diluted in PBS 10-fold at each transfer in 180 µL of sterile water and plated on YPD plates containing 100 µg/mL of Chloramphenicol to selectively grow yeast colonies and count the yeast population abundance after 3 days of incubation at 30 °C. Control plates for PBS contamination were plated following the same protocol but using only PBS. Then 100 µL of cell suspension were fixed in 4% Formaldehyde for 30 minutes in ice and then resuspended in PBS 1X overnight until sample staining for cytometry. Finally, from the same solution 800 µL of samples were transferred to 1.5 mL Eppendorf tubes and centrifuged at 10000 rpm for 10 minutes in a benchtop centrifuge. The supernatant was collected independently, and the pellet was stored for later DNA extraction. The fixed samples for cytometry were centrifugated at 5000 rpm for 5 minutes and resuspended in 300 µL of fresh PBS with 1 µL of SYTO BC (ThermoFisher) and 3 µL of quantification beads. The samples were acquired with a BDFortessa cytometer using the 488 nm laser and the YG-A filter at low speed. Acquired samples were then analysed using FlowJO™ and beads gated were used to calculate the relative proportion of cells in each sample.

### DNA extraction and 16S amplicon sequencing and analysis

DNA was extracted from the pellet recovered from the bee guts using the ZymoBIOMICS DNA Kits Miniprep from ZymoResearch™ (Ref D4300) following the manufacturer protocol. The DNA was sent for 16S amplicon sequencing at Eurofins Genomics. The V3-V4 region was amplified and generated fastq were then used for later analysis. The quality of the reads was assessed by plotting the quality profile along the read’s length with the function “plotQualityProfile” of the package dada2 (v1.26.0). Reads were trimmed to remove low quality regions using the function “filterAndTrim” cutting at 260 and 210 base pairs respectively the forward and reverse reads. The reads were then merged and the bimeras were then removed and the remaining reads mapped with the function “assign.Taxonomy” of the package dada2 to the Silva database (v138.1). Genus with a number of supporting reads below 20 were excluded. Genus’s abundance was then plotted using R and ggplot2.

### Data availability

Genomic data generated in this study are submitted to the NCBI SRA archive with the submission id SUB15827871. Proteomic data are included in the supplementary materials.

## Acknowledgements

We would like to thank Sonja Blasche for providing yeast strains, Alan Porter (IRAC coordinator) for providing information on insecticides mode of action, Mike Howell (High Throughput Screening STP, The Francis Crick Institute) for advice on compound library arrangement, Arianna Castorino for providing graphical illustrations. SM was funded by an HFSP long-term fellowship LT0018/2023-L DOI: 10.52044/HFSP.LT00182023. DS acknowledge funding by the Swiss National Science Foundation Grant (P500PB 211100), DB was supported by The Italian Ministry of Universities and Research (MUR) (Protocol No 2022LJN3LE; Cup: B53D23012130006), FDC was funded by National Recovery and Resilience Plan (NRRP), Mission 4 “Education and Research” Component M4C1. This project has received funding from the European Research Council under the European Union’s Horizon 2020 research and innovation programme (grant no. 866028 to K.R.P.), and from the UK Medical Research Council (project no. MC_PC_24010 to K.R.P.)

## Author contributions

SM and KP conceived the overall study. DB designed and planned the bee experiments. SM, FDC, SK, DS, AL, JK, SKP and LF performed the experiments, SM, SK and XJ performed data analysis. SM wrote the original manuscript draft with KP. All the authors reviewed the final manuscript.

## Competing Interest Statement

The authors declare no competing interests.

## Supplementary material

### List of supplementary tables

All tables are available to download from the submission system:

Supplementary Table 1 – Overview of microbial strains used in the study

Supplementary Table 2 – Compounds used for the adaptive laboratory evolution of *S. bombicola*

Supplementary Table 3 – Variants detected in the ALE experiment Supplementary Table 4 – Absolute fluorescent reading of ROS production Supplementary Table 5 – Absolute fluorescent reading of Mitochondrial biomass Supplementary Table 6 – Proteomic data of Paclobutrazol exposure

Supplementary Table 7 – Annotated proteins upregulated in the ancestral strain upon Paclobutrazol exposure

Supplementary Table 8 - Annotated proteins upregulated in the evolved strain upon Paclobutrazol exposure compared to the ancestral

Supplementary Table 9 – Habituation experiment in honeybees upon 3 doses treatment of Paclobutrazol

Supplementary Table 10 – Sucrose response experiment in honeybees upon 3 doses treatment of Paclobutrazol

Supplementary Table 11 – Habituation experiment in honeybees upon yeast supplementation and pesticide treatment

Supplementary Table 12 – Sucrose response experiment in honeybees upon yeast supplementation and pesticide treatment

Supplementary Table 13 – Bee gut yeast population and FACS population total estimates Supplementary Table 14 – Optical density recorded during the ALE experiment

Supplementary Table 15 – Compounds used in the initial screening and hits called for reduced growth

Supplementary Table 16 – 16S sequencing analysis results

---

**Supplementary Figure 1.**
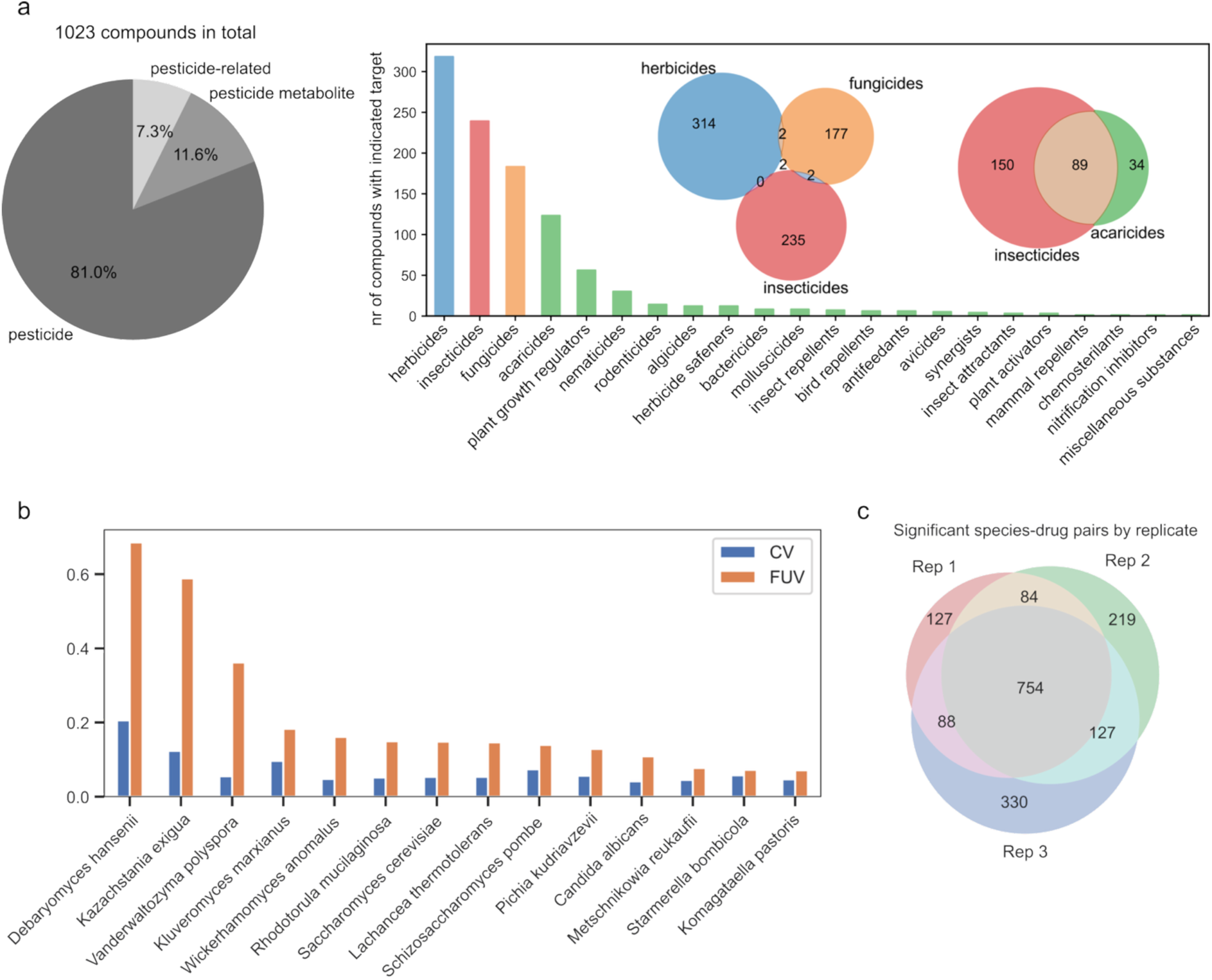
**a.** Pie chart of the compounds present in the library (left) and bar plot of compounds breakdown by classes. **b.** Coefficient of variation (CV) and fraction of unexplained variance (FUV) of the screened yeast species. **c.** Venn-diagram representing the matching hits across the three replicates growth experiments.

**Supplementary Figure 2.**
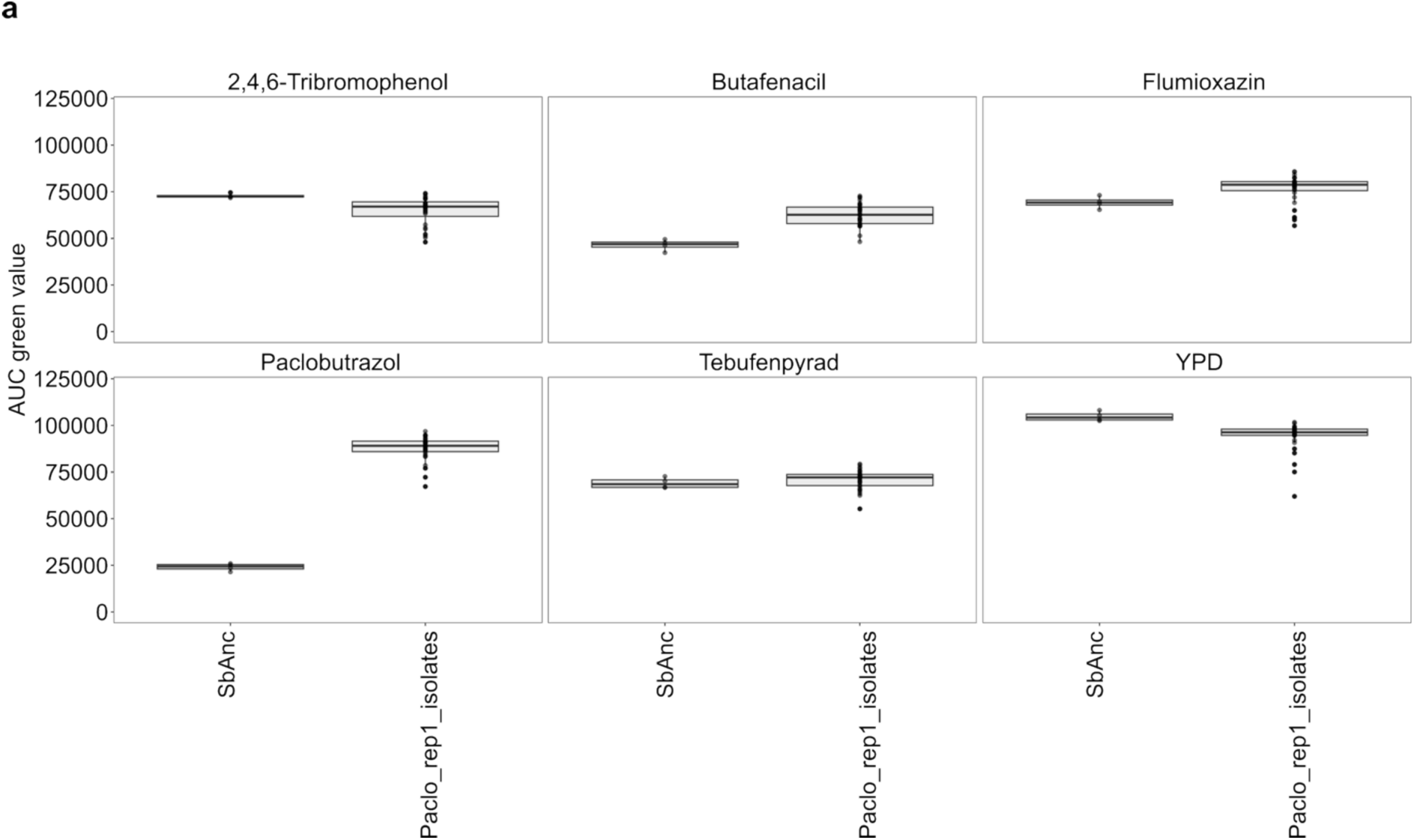
**a.** Area under the curve (AUC) measured as green value (proxy of optical density OD) of the evolved isolates from one selected population evolved in Paclobutrazol versus ancestral isolates (SbAnc). Each dot (n=24) is a single replicate. The concentrations tested are the same used for the ALE experiments. The evolved isolates showed an increased AUC in Paclobutrazol, Butafenacil and Flumioxazin compared to the ancestral (t-test two tail, p-value < 0.05)

**Supplementary Figure 3.**
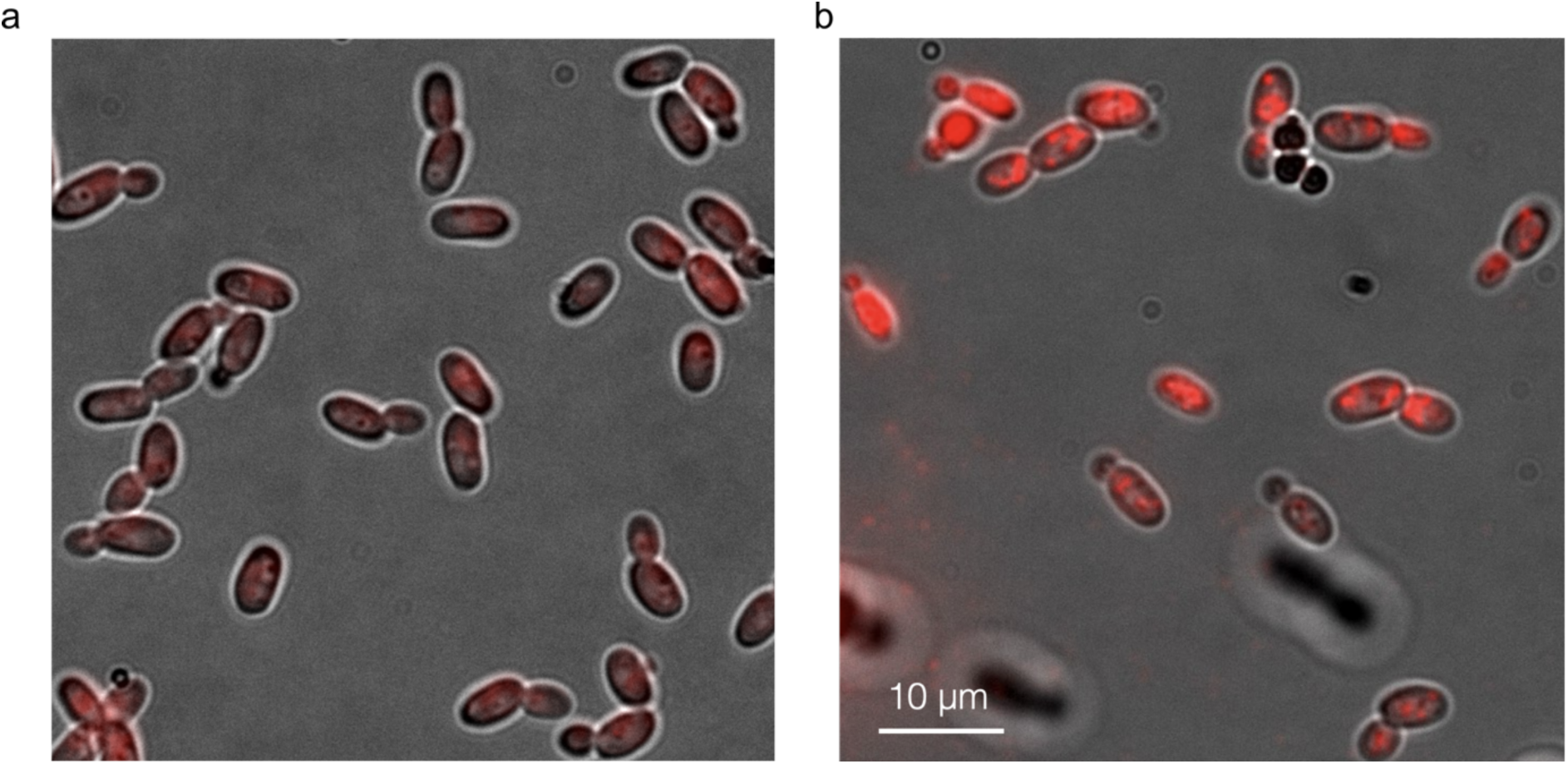
**a.** Yeast suspension of untreated (+DSMO) stained with Red MitoTracker for 15 minutes **b.** Yeast suspension of treated (+Paclobutrazol) stained with Red MitoTracker for 15 minutes. Scale bar is 10 µm.

**Supplementary Figure 4.**
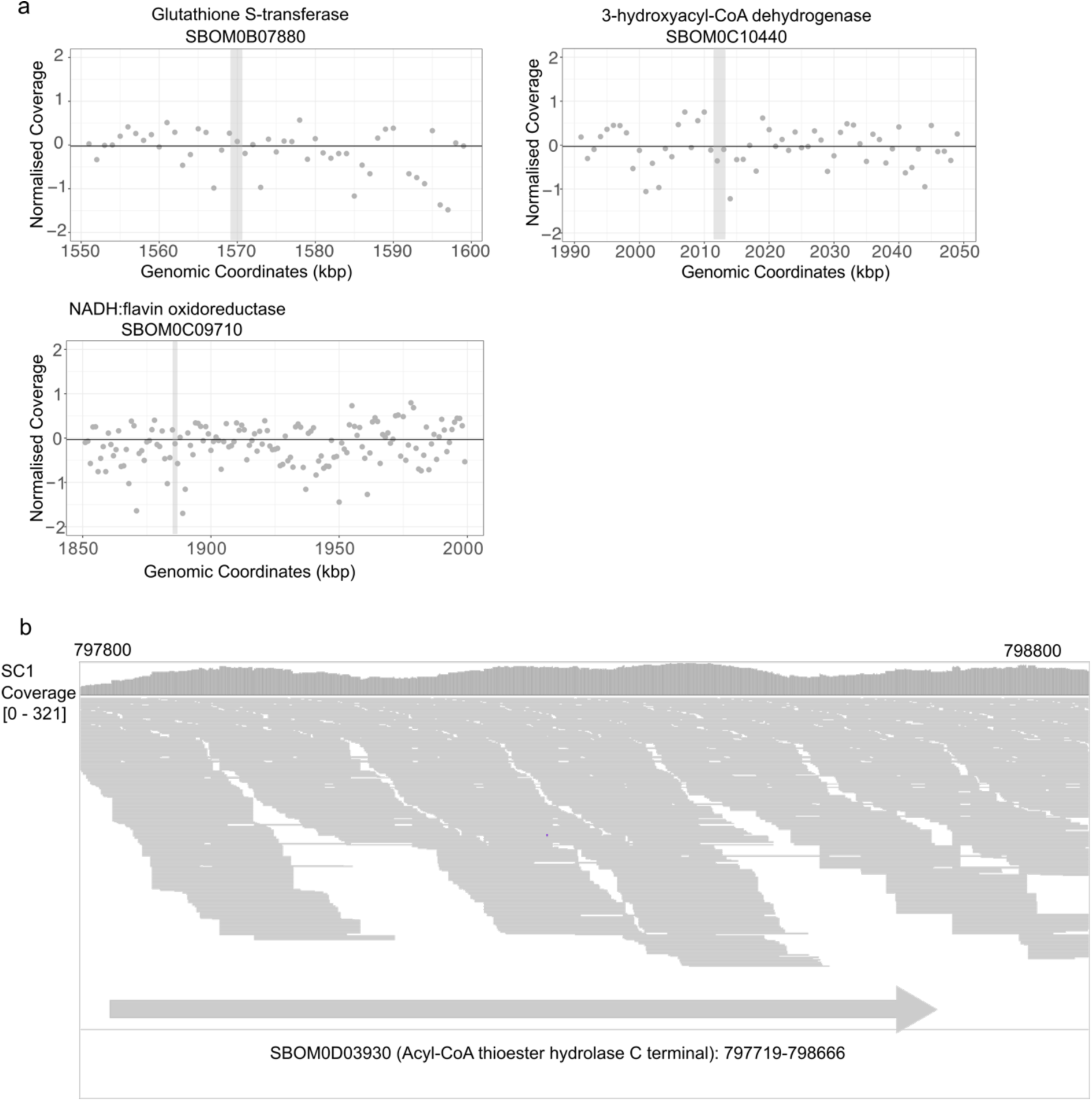
**a.** Normalised coverage across genomic regions bearing the HGT from bacteria. The grey bar highlights the region where the gene is found. The normalised coverage is calculated as the coverage per window of 1 kb divided by median chromosome coverage **b.** IGV condensed view of the reads on the region harbouring one HGT. The regions around the start and end of the gene do not display any lack in coverage of mis-mapping mate reads, indicating a true genomic insertion.

**Supplementary Figure 5.**
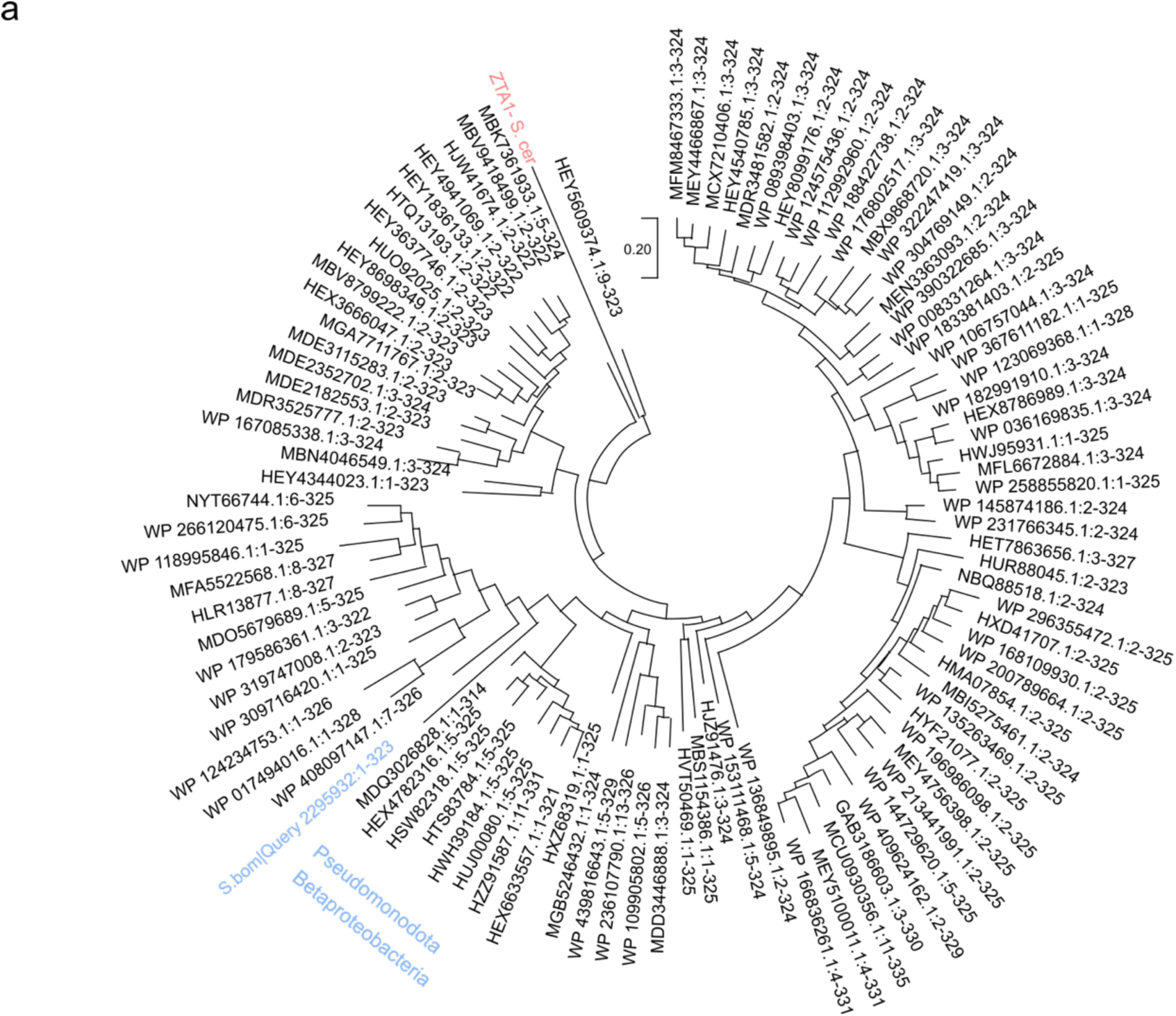
**a.** Phylogenetic relationship of the NADPH-quinone oxidase of *S. bombicola* (blue) relative to the top 100 hits on NCBI protein blast. The *saccharomyces* gene highlighted in red is added as outgroup

**Supplementary Figure 6.**
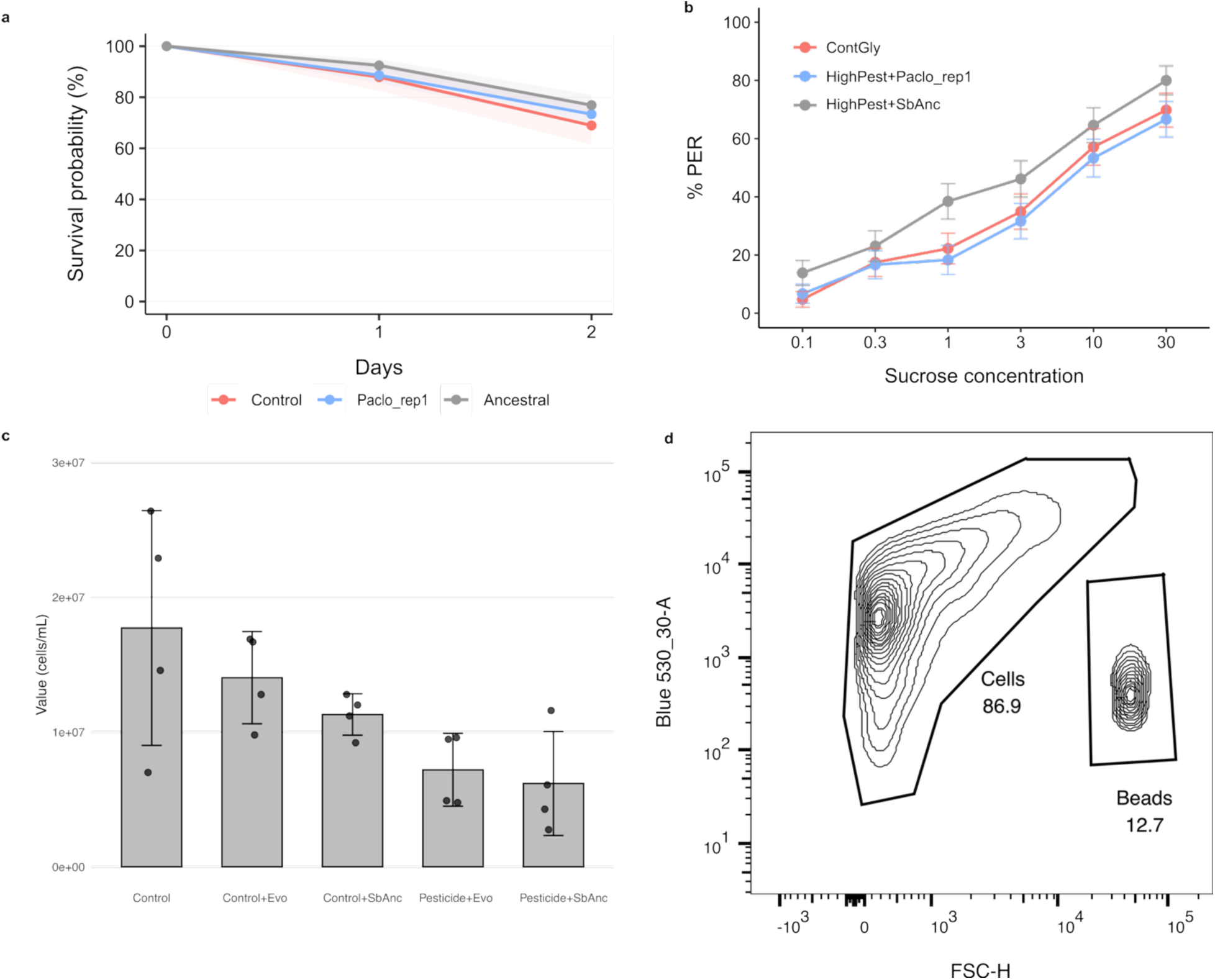
**a.** Survival proportion of treated bees with control, ancestral and evolved *S.* bombicola The y-axis report the percentage of survival at the different days of the treatment, day 1 and day 2. **b.** Sucrose responsiveness curve for the control and treated bees **c.** Microbial population density as estimated by FACS counting at the cytometry. Each dot is an individual gut sample (n=4), and the error bar represent the standard deviation. Population declined significantly only comparing the control to the treated samples (Pesticide+Evo p-value=0.041 or Pesticide+SbAnc, p-value=0.03) **c.** Gating strategy used for bacteria counting highlighting the two gates used after selecting events on the Blue 530-30 gate.

**Supplementary Figure 7.**
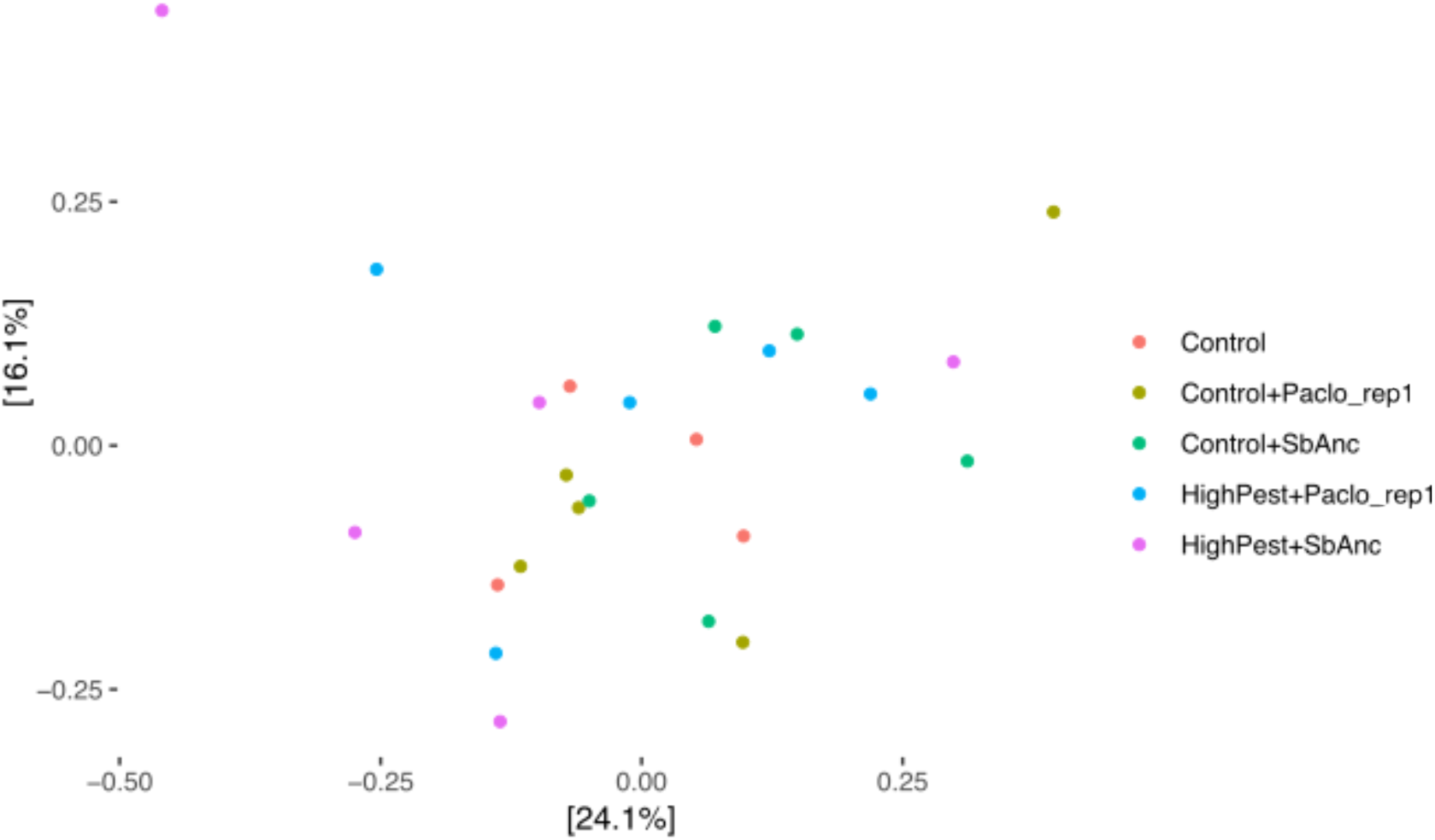
**a.** PCoA plot of the of Jaccard distances. X and Y axes represent PCA1 and PCA2. Jaccard distances were only statistical significant based on species threshold: Pairwise Permanova test on Jaccard distance with Bonferroni correction: Control versus Anc+High Conc: p-value<0.05) removing species with abundance lower than 0.1% did not show significant differences between groups (pairwise Permanova test on Jaccard distance with Bonferroni correction: Control versus Anc+High Conc: p-value>0.05).

**Supplementary Figure 8.**
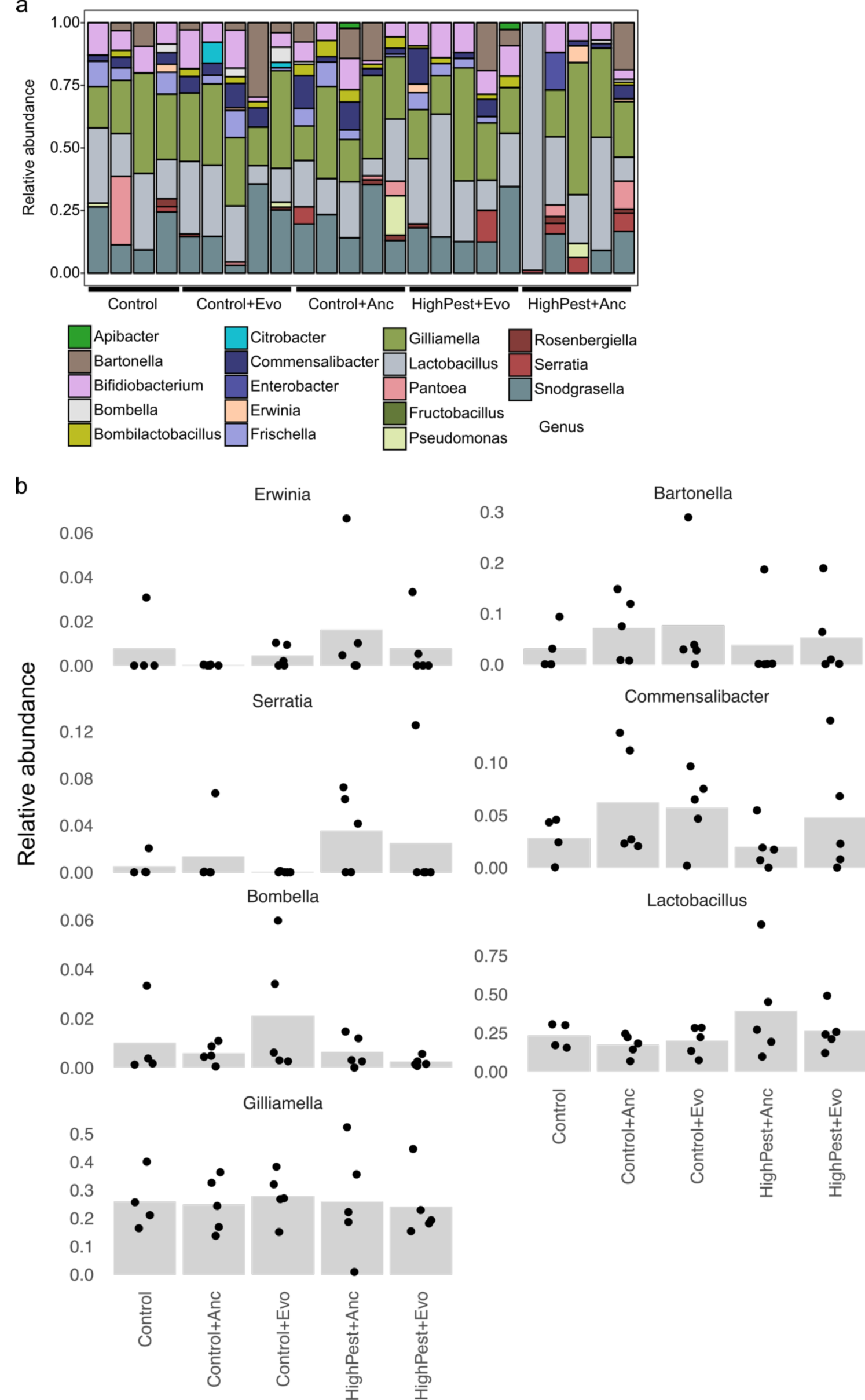
**a.** Relative abundances estimated by 16S sequencing of the bacterial composition of dissected bee guts (Control n=4 Treatments n=5). **b.** Bar plots representing core and accessory species of bacteria identified in the treated guts. The y-axis represents the relative abundance in each plot.

## Notes

### Competing Interest Statement

The authors have declared no competing interest.

